# Astrocytic control of extra-cellular GABA drives circadian time-keeping in the suprachiasmatic nucleus

**DOI:** 10.1101/2023.01.16.523253

**Authors:** A. P. Patton, E.L. Morris, D. McManus, H. Wang, Y. Li, J.W. Chin, M. H. Hastings

## Abstract

The hypothalamic suprachiasmatic nucleus (SCN) is the master mammalian circadian clock. Its cell-autonomous timing mechanism, a transcriptional/translational feedback loop (TTFL), drives daily peaks of neuronal electrical activity. Intercellular signals synchronize and amplify TTFL and electrical rhythms across the circuit. SCN neurons are GABAergic, but the role of GABA in circuit-level time-keeping is unclear. SCN slices expressing the GABA sensor iGABASnFR demonstrate a circadian oscillation of extracellular GABA ([GABA]_e_) that, counter-intuitively, runs in antiphase to neuronal activity, peaking in circadian night. Resolving this paradox, we found that [GABA]_e_ is regulated by GABA transporters (GATs), uptake peaking during circadian day. This is mediated by the circadian-regulated, astrocytically expressed GAT3 (*Slc6a11*). Clearance of [GABA]_e_ in circadian day facilitates neuronal firing, neuropeptide release and TTFL rhythmicity. Moreover, genetic complementation demonstrated that the astrocytic TTFL can alone drive [GABA]_e_ rhythms. Thus, astrocytic clocks maintain SCN circadian time-keeping by temporally controlling GABAergic inhibition of SCN neurons.

## Introduction

The suprachiasmatic nucleus (SCN) of the hypothalamus is the master circadian (~daily) clock in mammals. It coordinates the timing of distributed peripheral tissue clocks, thereby controlling adaptive behavioural and physiological rhythms. It is synchronised to environmental time by innervation from the retinohypothalamic tract (RHT) (Patton and Hastings, 2018), but the RHT is dispensable for robust SCN timekeeping. When isolated *ex vivo*, the SCN sustains precise circadian rhythms of molecular, electrical and metabolic activity.

At the cellular level, timekeeping pivots around a core transcriptional-translational feedback loop (TTFL) whereby the transcription factors CLOCK and BMAL1 drive expression of *Period* and *Cryptochrome* genes, and their encoded proteins, PER1/2, and CRY1/2, subsequently repress that transactivation. This generates a self-sustaining cycle of TTFL activity and its dependent transcriptional outputs (Cox and Takahashi, 2019), including ion channels (Harvey et al., 2020). These in turn direct daily rhythms in SCN neuron electrical excitability (Colwell, 2011), peaking in circadian day, with quiescence during circadian night (Brancaccio et al., 2013; Brancaccio et al., 2017; Colwell, 2011).

Cell-autonomous time-keeping is augmented by reciprocal circuit-level neuron-to-glia signalling that determines emergent properties of the SCN network: precise high amplitude oscillation, tightly defined ensemble period and phase, and spatiotemporally complex cellular synchrony (Hastings et al., 2018; Patton et al., 2020). These networklevel mechanisms are so powerful they sustain robust rhythms with periods ranging between 16 and 42 hours in various genetic/pharmacological contexts (Patton et al., 2016). Neuronal electrical rhythms are integral to this synchronisation through appropriately timed neurotransmitter/neuropeptide release: pharmacological blockade of neuronal firing (Yamaguchi et al., 2003) and synaptic vesicle release (Ferrari et al., 2011) desynchronise and damp the network.

SCN neurons are neurochemically heterogeneous due to their complex expression of a wide array of neuropeptides and receptors (Abrahamson and Moore, 2001; Morris et al., 2021; Park et al., 2016; Wen et al., 2020; Xu et al., 2021), but are also homogeneous insofar as almost all SCN neurons utilise the inhibitory small neurotransmitter γ-aminobutyric acid (GABA) (Albers et al., 2017). GABAergic neurotransmission is mediated via ionotropic GABA_A_ or metabotropic GABA_B_-receptors, and both subtypes are expressed within the SCN (Albers *et al*., 2017) together with GABA transporters (GATs) (Moldavan et al., 2015). Nevertheless, little is known about the role of GABA in steady-state network timekeeping (Albers *et al*., 2017; Ono et al., 2021). It has been considered as a counterbalance to neuropeptidergic signalling (Freeman et al., 2013a), a synchronising factor (Liu and Reppert, 2000), a modulator of encoded phase (Albers *et al*., 2017) and a network adaptor to seasonal changes in day length (Farajnia et al., 2014; Myung et al., 2015; Rohr et al., 2019). Pharmacological and genetic disruption of GABAergic signalling does not, however, disrupt TTFLoscillations and circadian firing of SCN *ex vivo* (Freeman et al., 2013b; Ono et al., 2019; Patton *et al*., 2016). This suggests that although GABA is important *in vivo* as an output from the SCN acting on distal targets (via GABAergic terminals) (Maejima et al., 2021; Ono *et al*., 2019) it is largely dispensable at the SCN network level. Indeed, how could an exclusively GABAergic circuit sustain circadian electrical activity, when increased neuronal firing would become inhibitory to the network?

To reconcile this paradox and identify a role for GABA within the SCN network, we first monitored real-time extracellular GABA dynamics ([GABA]_e_) in free-running SCN explants. This revealed a counter-intuitive daily bulk [GABA]_e_ flux in antiphase to (GABAergic) neuronal activity. This rhythm is not, however, driven by circadian variation in neuronal activity, but is rather driven by day-time GABA uptake via GATs, GAT1 (*Slc6a1*, mGAT1) and GAT3 (*Slc6a11*, mGAT4). Furthermore, this rhythmic uptake is controlled by the cell-autonomous circadian clock of astrocytes. This reveals a novel GABAergic network mechanism whereby active daytime uptake is permissive to neuronal activation and neuropeptide release during circadian day. Thus, the SCN circuit has devolved GABAergic control of its neurons to its astrocytes and their cell-autonomous clock to sustain and encode circadian time.

## Results

### Extracellular GABA is rhythmic in the SCN and peaks during circadian night

To determine the circadian dynamics of [GABA]_e_, we transduced SCN explants with adeno-associated viral vectors (AAVs) encoding the fluorescent GABA reporter iGABASnFR (Marvin et al., 2019) controlled by the neuronal *synapsin* (*Syn*) promoter. Appropriate membrane-targeted fluorescence (Supplementary Figure 1) was observed across the explant one-week post-transduction (Figure 1A). Fluorescence was recorded for ~5 days in tandem with a bioluminescent TTFL reporter, PER2::Luciferase (PER2::LUC), which provided a circadian phase reference: PER2::LUC peaks at circadian time (CT)12, the start of circadian night (Figure 1A). Imaging revealed rhythms in [GABA]_e_ with a robust waveform consisting of a broad, flat peak and a sharp trough. The circadian properties of the [GABA]_e_ rhythm were comparable with those of PER2::LUC, with identical periods (Figure 1B, period: PER2::LUC 24.14±0.12h vs [GABA]_e_ 24.03±0.12h, paired two-tailed t-test t(7)=1.23, p=0.26), rhythm quality (Figure 1C, rhythmicity index (RI): PER2::LUC 0.54±0.01 vs [GABA]_e_ 0.52±0.03, paired two-tailed t-test t(7)=1.45, p=0.19) and precision (Figure 1D, relative amplitude error (RAE): PER2::LUC 0.06±0.01 vs [GABA]_e_ 0.08±0.01, paired two-tailed t-test t(7)=1.80, p=0.12). Intriguingly, however, [GABA]_e_ peaked during circadian night in all slices (CT19.7±0.48, Rayleigh test p<0.0001, R=0.95) (Figure 1E), a counter-intuitive finding given that the exclusively GABAergic SCN neurons are electrically active (and releasing synaptic GABA) in circadian daytime (~CT06).

**Figure 1:**
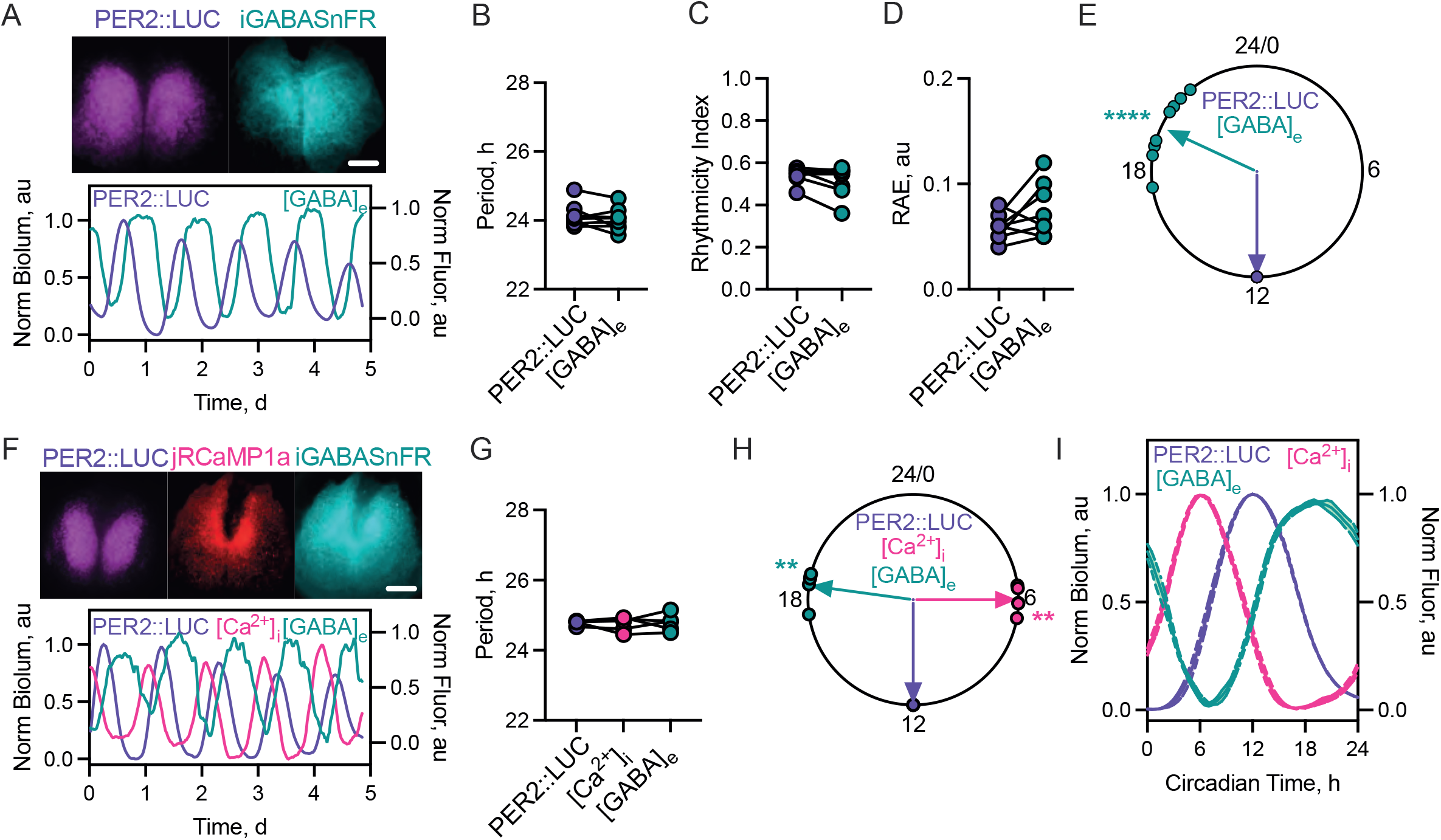
iGABASnFR reports robust oscillations in [GABA]_e_ in antiphase to SCN neuronal activity. A. (Upper) Average Z-projections of PER2::LUC (left) bioluminescence and *Syn*.iGABASnFR (right) fluorescence in an *ex vivo* SCN slice. (Lower) Example normalised PER2::LUC bioluminescence and *Syn*.iGABASnFR fluorescence ([GABA]_e_). B-D. Histograms showing aggregate statistics for PER2::LUC and [GABA]_e_ rhythms paired by SCN slice for: period (B), rhythmicity index (C) and RAE (D). E. Circular Rayleigh plot showing the relative phasing of peak [GABA]_e_ aligned on a slice-by-slice basis to peak PER2::LUC. F. (Upper) Average Z-projections of PER2::LUC (left) bioluminescence, *Syn*.jRCaMP1a (middle) and *Syn*.iGABASnFR (right) fluorescence. (Lower) Example trace showing normalised multiplexed PER2::LUC, *Syn*.jRCaMP1a ([Ca^2+^]_i_) and *Syn*.iGABASnFR ([GABA]_e_). G. Histogram showing aggregate period for PER2::LUC, [Ca^2+^]_i_ and [GABA]_e_ rhythms paired by SCN slice. H. Circular Rayleigh plot showing the relative phasing of the peak in [Ca^2+^]_i_ and [GABA]_e_ aligned to peak PER2::LUC. I. Normalised single cycles of PER2::LUC, [Ca^2+^]_i_ and [GABA]_e_ rhythms aligned to peak PER2::LUC. In all plots, points represent SCN with paired measures joined. Lines/shading represent mean±SEM. For B-E N=8, G-I N=4. Scale bar=250μm.

Due to this unexpected observation, we then co-registered [GABA]_e_ explicitly with neuronal activity, reflected by intracellular calcium levels ([Ca^2+^]_i_). PER2::LUC SCN explants were co-transduced with *Syn*.iGABASnFR and *Syn*.jRCaMP1a. This, again, revealed robust rhythms of [GABA]_e_ alongside circadian oscillations of neuronal [Ca^2+^]_i_ (Figure 1F). All three circadian markers had identical periods (Figure 1G, PER2::LUC 24.79±0.03h vs [Ca^2+^]_i_ 24.74±0.09h vs [GABA]_e_ 24.78±0.11h, repeated-measures one-way ANOVA, F(2, 8)=0.11, p=0.90). As anticipated, [Ca^2+^]_i_ peaked in the day (CT5.99±0.22) whereas in the same slices [GABA]_e_ again peaked in circadian night (CT18.51±0.27h), significantly delayed relative to [GABA]_e_ (paired two-tailed t-test, t(4)=34.51, p<0.0001) across all slices tested (Rayleigh test: [Ca^2+^]_i_ p=0.001, R=0.99; [GABA]_e_ p=0.001, R=0.99) (Figure 1H). Notably, average, single-cycle alignment of [Ca^2+^]_i_ and [GABA]_e_ rhythm waveforms mirrored one another: [GABA]_e_ dynamics appearing as an inversion of those of neuronal [Ca^2+^]_i_ (Figure 1I). In SCN explants, therefore, the circadian rhythm of [GABA]_e_ sits in antiphase to neuronal rhythmicity, peaking in circadian night.

### Reduced GABAergic tone during the day facilitates network timekeeping

To explore the functional consequences of the [GABA]_e_ rhythm, we artificially clamped GABAergic tone into a chronic “high” condition, mimicking night-time, by using the GABA_A_- and GABA_B_-receptor specific agonists muscimol and (R)-baclofen, respectively. Chronic activation of GABA_A_-receptors by muscimol (Figure 2A) dose-dependently lengthened circadian period (Figure 2B, one-way ANOVA F(4, 24)=13.91, p<0.0001), and reduced both precision (RAE) (Figure 2C, one-way ANOVA F(4, 24)=12.21, p<0.0001) and amplitude (Figure 2D, one-way ANOVA F(4, 24)=140.8, p<0.0001) without lasting effects on the oscillation, which was restored following wash-out (Supplementary Figure 2A). This reduction in amplitude was associated with a marked suppression of both the level of the circadian trough (Supplementary Figure 3A, one-way ANOVA F(4, 24)=32.52, p<0.0001) and peak (Supplementary Figure 3A, one-way ANOVA F(4, 24)=124.3, p<0.0001), consistent with an electrically suppressed SCN (Colwell, 2011). Intriguingly, the trough appeared more sensitive to muscimol at lower concentrations than did the peak (Supplementary Figure 3A). In contrast, chronic activation of GABA_B_ receptors by (R)-baclofen (Figure 2E) did not alter TTFL period (Figure 2F, one-way ANOVA F(4, 20)=2.39, p=0.09), precision (Figure 2G, one-way ANOVA F(4, 20)=1.647, p=0.20) or amplitude (Figure 2H, one-way ANOVA F(4, 20)=1.54, p=0.23). Although there was no effect on overall amplitude, chronic activation of GABA_B_-receptors slightly suppressed the trough (Supplementary Figure 3B, one-way ANOVA F(4, 20)=3.252, p=0.0330) without affecting the peak (Supplementary Figure 3B, one-way ANOVA F(4, 20)=2.50, p=0.07). Thus, whereas sustained activation of GABA_A_-receptors compromised the TTFL, the TTFL was not affected by sustained activation of GABA_B_-receptors.

**Figure 2:**
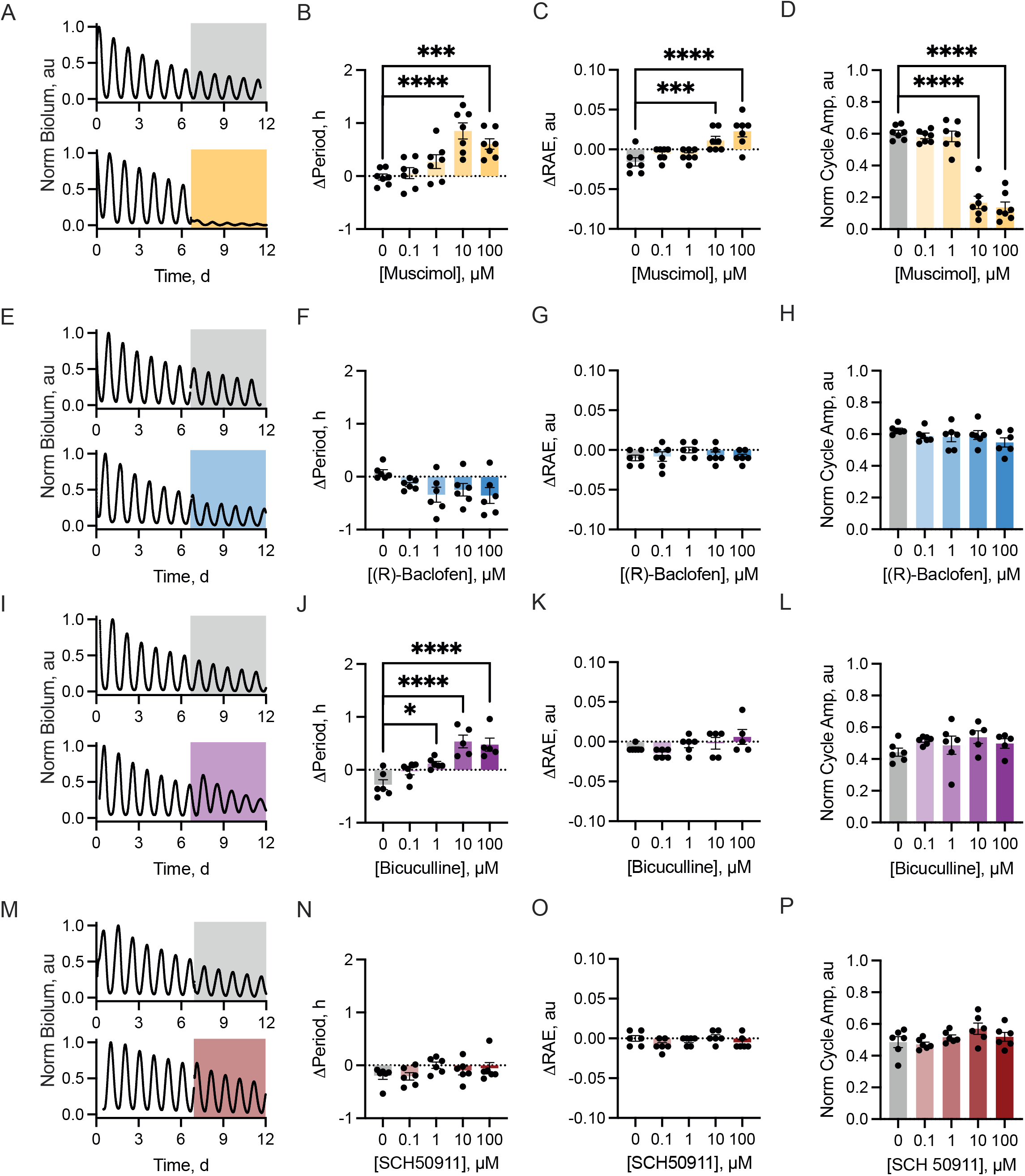
Chronic activation or inactivation of GABA_A_ receptors alters SCN network properties. A. PER2::LUC bioluminescence from SCN treated with vehicle (upper, grey) or 100μM muscimol (lower, orange). B-D. Histogram showing the change in period (B), RAE (C) and normalised amplitude (D) between baseline and treatment intervals versus muscimol concentration. E. PER2::LUC bioluminescence from SCN treated with vehicle (upper, grey) or 100μM (R)-baclofen (lower, blue). F-H. Histogram showing the change in period (F), RAE (G) and normalised amplitude (H) between baseline and treatment intervals versus (R)-baclofen concentration. I. PER2::LUC bioluminescence from SCN treated with vehicle (upper, grey) or 100μM (+)-bicuculline (lower, purple). J-L. Histogram showing the change in period (J), RAE (K) and normalised amplitude (L) between baseline and treatment intervals versus (+)-bicuculline concentration. M. PER2::LUC bioluminescence from SCN treated with vehicle (upper, grey) or 100μM SCH50911 (lower, maroon). N-P. Histogram showing the change in period (N), RAE (O) and normalised amplitude (P) between baseline and treatment intervals versus SCH50911 concentration. In all plots, points represent SCN (N≥5 for all concentrations) and bars represent mean±SEM.

To complement these observations on high GABA tone, we then clamped GABAergic signalling into a chronic “low” condition to mimic daytime by using the GABA_A_- or GABA_B_-receptor specific antagonists (+)-bicuculline and SCH50911, respectively. Blockade of GABA_A_-receptors with (+)-bicuculline (Figure 2I) slightly lengthened period at high doses (Figure 2J, one-way ANOVA F(4,23)=14.16, p<0.0001) without altering precision (Figure 2K, one-way ANOVA F(4,23)=2.15, p=0.11) or amplitude (Figure 2L, one-way ANOVA F(4,23)=0.96, p=0.45) and without lasting effects on the ongoing oscillation, which continued following wash-out (Supplementary Figure 2B). Despite the lack of effect of (+)-bicuculline on the overall cycle amplitude, there was a dose-dependent increase in the level of both the circadian trough (Supplementary Figure 3C, one-way ANOVA F(4,23)=3.79, p=0.02) and peak (Supplementary Figure 3C, one-way ANOVA F(4,23)=8.65, p=0.0002), consistent with increased electrical excitability feeding into the TTFL (Colwell, 2011). In contrast, increasing doses of the GABA_B_-receptor antagonist SCH50911 (Figure 2M) did not alter period (Figure 2N, one-way ANOVA F(4,25)=1.32, p=0.29), precision (Figure 2O, one-way ANOVA F(4,25)=1.64, p=0.20) or amplitude (Figure 2P, one-way ANOVA F(4,25)=1.99, p=0.13). Equally, SCH50911 had no effect on the trough (Supplementary Figure 3D, one-way ANOVA F(4,25)=1.62, p=0.20) or peak (Supplementary Figure 3D, one-way ANOVA F(4,25)=1.35, p=0.28). Thus, loss of GABA_A_-signalling affects TTFL period and range of operation, whereas loss of GABA_B_ does not. Taken together, the data from agonists and antagonists demonstrate that GABAergic signalling acts via the GABA_A_- but not GABA_B_-receptors to suppress TTFL function in the SCN. This suggests that the low level of GABA during circadian day facilitates day-time neuronal activity, which in turn boosts SCN network function (Colwell, 2011).

### GABA transporters facilitate ongoing SCN rhythmicity

To reconcile the apparently paradoxical observation that neuronal activity (which should drive GABA release) occurs when [GABA]_e_ is lowest and [GABA]_e_ is highest at night when neurons are inactive, we investigated mechanisms that could control GABA flux. First, we explored published single-cell RNA-sequencing (scRNA-seq) data from SCN explants harvested during circadian day (CT7.5) or night (CT15.5) (Morris et al, 2021) to determine the expression of genes encoding the four canonical GATs: GAT1 (*Slc6a1*, mGAT1), GAT2 (*Slc6a13*, mGAT3), GAT3 (*Slc6a11*, mGAT4) and BGT1 (*Slc6a12*, mGAT2). Data were clustered irrespectively of time of day into three principal cell groups: neurons (defined by *Slc32a1, Tubb3* and *Celf4* expression), astrocytes (defined by *Aldh1l1, Gfap* and *Aqp4* expression) and other (defined by exclusion of these markers). Cell-clusters were then divided by time-of-day, and relative expression levels of the four *Gat* genes evaluated. Genes encoding betaTubulin-3 (*Tubb3*) and ALDH1L1 (*Aldh1l1*) were used as internal controls to confirm specific segregation of neuronal and astrocyte populations (Figure 3A). This revealed *Gat1* and *Gat3* as the principal GATs within *ex vivo* SCN (with little to no *Gat2* or *Bgt1* expression), and expression of GAT1 and GAT3 across the SCN was confirmed by immunostaining (Supplementary Figure 4A). scRNAseq revealed that *Gat1* was expressed by SCN astrocytes and neurons, while *Gat3* was specifically enriched in SCN astrocytes (Figure 3A) (Barca-Mayo et al., 2017; Coomans et al., 2021; Moldavan *et al*., 2015). Furthermore, testing for differential expression in astrocytes between the day and night revealed a significant temporal variation in the levels of *Gat1* and *Gat3*, with expression being higher during the circadian day (Figure 3B). This was confirmed by qPCR (Supplementary Figure 4B). The fact that the transcriptomic data predict that GATs are present at a higher abundance during the circadian day when [GABA]_e_ is low and at a lower abundance when [GABA]_e_ is high highlights them as candidates to generate the observed dynamics in [GABA]_e_.

**Figure 3:**
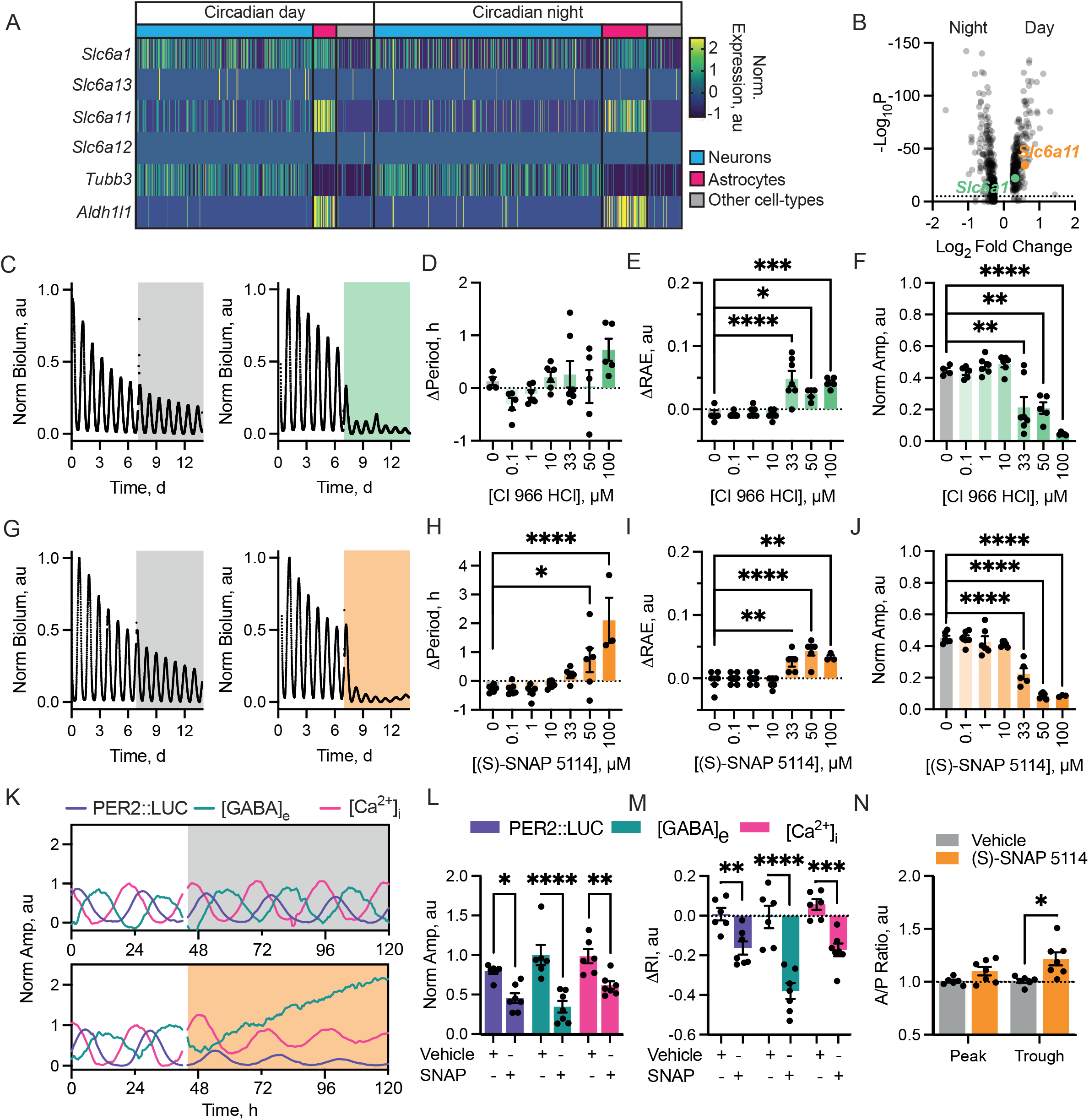
Inhibition of GATs disrupts circadian timekeeping and compromises [GABA]_e_ dynamics. A. Heatmap of normalised expression levels of *Slc6a1* (GAT1, mGAT1), *Slc6a13* (GAT2, mGAT3), *Slc6a11* (GAT3, mGAT4) and *Slc6a12* (BGT1, mGAT2) alongside the neuronal *Tubb3* and astrocytic *Aldh1l1* markers. Cells-types are indicated by coloured bars: neurons (blue), astrocytes (magenta) and all others (grey). Time of day at which cells were harvested is indicated. Re-analysed from (Morris *et al*., 2021). B. Volcano plot of differential day-night gene expression in astrocytes. Horizontal line indicates the significance cut-off. *Slc6a1* (GAT1, green) and *Slc6a11* (GAT3, orange) are significantly upregulated in day-time. C. PER2::LUC trace from SCN treated with vehicle (left, grey) or 50μM CI 966 HCl (right, green). D-F. Histogram showing change in period (D), relative amplitude error (RAE) (E) and normalised amplitude (F) between baseline and treatment intervals versus CI 966 HCl concentration. G. PER2::LUC from vehicle (left, grey) or 50μM (S)-SNAP 5114 (right, orange) treated slices. H-J. Histogram showing the change in period (H), RAE (I) and normalised amplitude (J) between baseline and treatment intervals versus (S)-SNAP 5114 concentration. K. Example PER2::LUC, [GABA]_e_ and neuronal [Ca^2+^]_i_ traces from vehicle (left, grey) or 50μM (S)-SNAP 5114 (SNAP) (right, orange) treated slices. L. Min-to-max amplitude of the treatment interval normalised to the baseline for vehicle or SNAP-treated SCN comparing effects on PER2::LUC, [GABA]_e_ and [Ca^2+^]_i_. M. Change in rhythmicity index (ΔRI) between baseline and treatment intervals of vehicle or SNAP-treated SCN for PER2::LUC, [GABA]_e_ and neuronal [Ca^2+^]_i_. N. Ratio of [GABA]_e_ predicted/actual peak or trough amplitudes during vehicle or SNAP-treatment. For D-F, N≥4; H-J, N≥3; L-N, N≥7. In all plots, points represent SCN and bars represent mean±SEM, except B where points represent genes.

To determine the contribution of these transporters to the SCN TTFL oscillation, following an initial baseline recording we treated PER2::LUC explants with the GAT1-specific inhibitor CI966-HCl or the GAT3-specific inhibitor (S)-SNAP 5114 (Figure 3C). Increasing dose of the GAT1 inhibitor CI966-HCl caused a small increase in period (Figure 3D, one-way ANOVA F(6,31)=2.71, p=0.03). This was accompanied by decreased precision (Figure 3E, one-way ANOVA F(6,31)=14.01, p<0.0001) and suppression of the amplitude (Figure 3F, one-way ANOVA F(6,38)=26.95, p<0.0001) with no lasting effect on the ongoing oscillation, which returned immediately following washout (Supplementary Figure 5A). These changes occurred at relatively high concentrations, between 33 and 100μM, however, which are ~100-times the IC_50_ for CI966-HCl at GAT1 (IC_50_ 0.1μM, (Borden et al., 1994b)). Furthermore, at 100μM CI966-HCl could also inhibit GAT3 transporters (IC_50_ for GAT3 300μM, (Borden *et al*., 1994b)). We therefore interpreted this suppression as a consequence of (at least partial) inhibition of GAT3 alongside GAT1.

To clarify this, we recorded PER2::LUC rhythms from explants treated with a range of doses of the GAT3-specific inhibitor (S)-SNAP 5114 (Figure 3G). Again, as dose increased, period lengthened (Figure 3H, one-way ANOVA F(7,36)=9.40, p<0.0001), precision decreased (Figure 3I, one-way ANOVA F(7,36)=14.01, p<0.0001) and amplitude was suppressed (Figure 3J, one-way ANOVA F(7,36)=56.66, p<0.0001). These effects were reversible, but in contrast to the immediate restoration following removal of CI966-HCl, the TTFL took several cycles to be fully restored: amplitude progressively expanded cycle-to-cycle post-washout (Supplementary Figure 5B, C). Importantly, the concentrations of (S)-SNAP 5114 that maximally produced these sustained effects, between 33 and 50μM, are within 6-10 times the IC_50_ for GAT3 transporters (IC_50_ 5μM, (Borden et al., 1994a)), thereby confirming the predominant role of GAT3 in sustaining TTFL function in the SCN.

To assess how treatment with (S)-SNAP 5114 alters the dynamics of [GABA]_e_, we recorded bioluminescent and fluorescent emissions from PER2::LUC SCN explants expressing *Syn*.iGABASnFr and *Syn*.jRCaMP1a, before and during treatment with either vehicle or 50μM (S)-SNAP 5114 (Figure 3K). Vehicle had no effect, but GAT3 inhibition rapidly elevated [GABA]_e_ and suppressed neuronal [Ca^2+^]_i_ rhythms and PER2::LUC (Figure 3K,L). Quantification of this suppression as the absolute peak-to-trough excursion was significant relative to vehicle treatment across all reporters (Figure 3L) (repeated-measures two-way ANOVA: Treatment-effect F(1,6)=61.17, p=0.0002; Reporter-effect F(2,12)=2.40, p=0.13; Interaction F(2,9)=3.19, p=0.09). Nevertheless, while rhythmicity was reduced across all reporters relative to vehicle (Figure 3M) (repeated-measures two-way ANOVA Treatment-effect F(1,6)=40.87, p=0.0007), the effect was not equal across reporters (Reporter-effect F(2,12)=5.69, p=0.018; Interaction F(2,9)=32.11, p<0.0001). Post-hoc multiple comparisons revealed that reduction by (S)-SNAP 5114 was most severe in the [GABA]_e_ oscillation (Sidak’s multiple comparisons test, DMSO vs (S)-SNAP 5114 ΔRI: PER2::LUC 0.01±0.03 vs −0.16±0.03 au, p=0.007; [GABA]_e_, 0.01±0.06 vs −0.38±0.04 au, p<0.0001; [Ca^2+^]_i_ 0.06±0.03 vs −0.17±0.03 au, p=0.0003) (Figure 3M). Chronic GAT3 inhibition therefore has potent effects on the ongoing TTFL oscillation through disruption of neuronal activity (as evidenced by [Ca^2+^]_i_ report), presumably through sustained elevation and severe disruption of the [GABA]_e_ rhythm.

GAT3 inhibition raised [GABA]_e_, reduced network rhythmicity and eliminated [GABA]_e_ rhythmicity in SCN explants. The scRNAseq data suggested that the [GABA]_e_ rhythm is generated by a day-time GAT-mediated uptake of GABA from the extracellular space, meaning that GAT activity should account for the trough but not peak of [GABA]_e_. To test this, we extrapolated the trough and peak levels from baseline recordings forward into treatment intervals (Supplementary Figure 6), enabling us to quantify the relative change in the peak and trough of the [GABA]_e_ rhythm whilst under treatment (Figure 3N). This revealed a significant change in these metrics under treatment with (S)-SNAP 5114, but not vehicle (repeated-measures two-way ANOVA: Treatment-effect F(1,6)=7.52, p=0.03) which was not equal between the peak and the trough of the rhythm (repeated-measures two-way ANOVA: Peak/trough-effect F(1,6)=10.13, p=0.019; Interaction, F(1,4)=9.02, p=0.040). The flattening of the [GABA]_e_ rhythm was caused by elevation of the circadian trough, but not the peak of the oscillation (Sidak’s multiple comparisons test Vehicle vs (S)-SNAP 5114 actual/predicted ratio: Peak 1.01±0.01 vs 1.10±0.04 au, p=0.25; Trough 1.01±0.02 vs 1.22±0.06 au, p=0.01) (Figure 3N). Thus, under free-running conditions, GAT3 in the SCN is responsible for taking up GABA released during the day, presumably to facilitate neuronal firing, which in turn is integral for high-amplitude, precise oscillation across the circuit.

### Daytime uptake of GABA permits neuropeptidergic signalling

Our model predicts that the daily [GABA]_e_ rhythm, driven principally by day-time up-regulated GAT3 activity, facilitates increased neuronal excitability in the presence of enhanced neuronal GABA release. In the SCN, neuropeptide release is the critical functional consequence of neuronal excitability in sustaining circuit-level oscillations. To interrogate the relationship between [GABA]_e_ and neuropeptide release, we recorded extracellular VIP levels ([VIP]_e_) via AAV-dependent expression of the fluorescent VIP sensor, GRAB_VIP1.0_ (*Syn*.GRAB_VIP1.0_). In this reporter, sensitivity to VIP is conferred by a catalytically dead VPAC2 receptor coupled to circularly permuted EGFP (Wang et al., 2022). We recorded *Syn*.GRAB_VIP1.0_ in SCN explants, phase-registering the signal against PER2::LUC oscillations (Figure 4A). This revealed pronounced [VIP]_e_ rhythms with properties comparable to the PER2::LUC rhythm (Figure 4B-D, PER2::LUC vs [VIP]_e_: period: 25.11±0.35h vs 25.21±0.39h, paired twotailed t-test t(6)=0.64, p=0.55; rhythmicity index: 0.49±0.06 vs 0.41±0.06, paired twotailed t-test t(6)=2.42, p=0.052; precision: 0.07±0.01 au vs 0.07±0.01, paired two-tailed t-test t(6)=0.42, p=0.69). Importantly, the rhythm in [VIP]_e_ peaked in mid-circadian day, CT6.97±0.5h (Figure 4E, Rayleigh test R=0.95, p<0.0001) consistent with previous reports (Ono et al., 2023).

**Figure 4:**
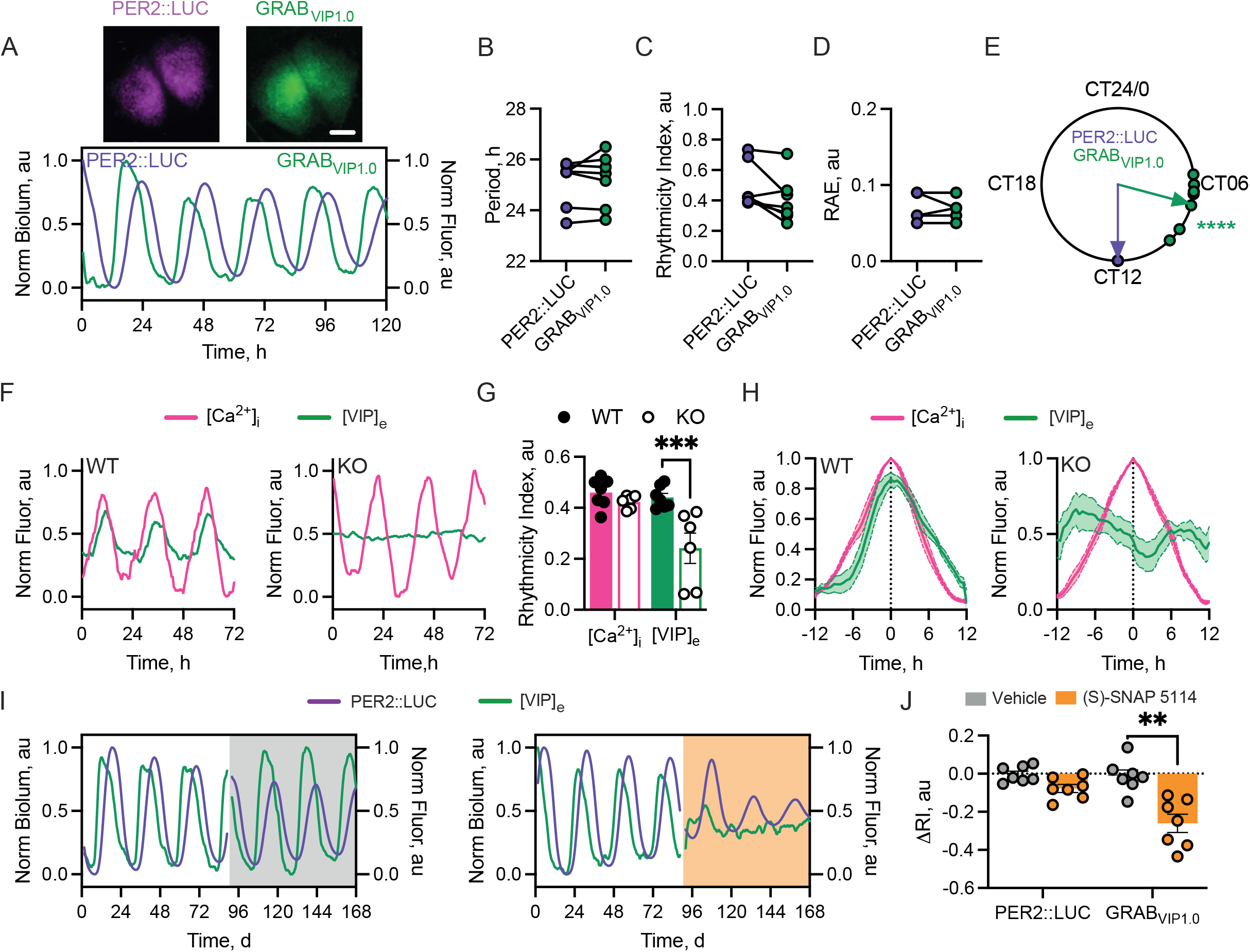
[VIP]_e_ in the SCN is rhythmic and is disrupted by GAT3 inhibition. A. (Upper) Average Z-projections of PER2::LUC (left) and GRAB_VIP1.0_ (right) in an *ex vivo* SCN slice. (Lower) Example normalised PER2::LUC and *Syn*.GRAB_VIP1.0_ fluorescence ([VIP]_e_). B-D. Histograms showing aggregate statistics for PER2::LUC and [VIP]_e_ rhythms for: period (B), rhythmicity index (C) and relative amplitude error (RAE) (D). E. Circular Rayleigh plot showing the relative phasing of the peak in [VIP]_e_ aligned on a slice-by-slice basis to peak PER2::LUC. F. Example normalised fluorescence traces showing [VIP]_e_ alongside neuronal [Ca^2+^]_i_ from VIP-WT (left) or VIP-KO (right) SCN. G. Histogram showing rhythmicity index of [VIP]_e_ and neuronal [Ca^2+^]_i_ in VIP-WT (filled circles/bars) or VIP-KO (open circles/bars) SCN. H. Average normalised [VIP]_e_/[Ca^2+^]_i_ single-cycles aligned by [Ca^2+^]_i_ for VIP-WT (left) and VIP-KO (right) SCN. I. Example PER2::LUC and [VIP]_e_ for vehicle (left, grey) or 50μM (S)-SNAP 5114 (SNAP) (right, orange) treated SCN. J. Change in rhythmicity index (ΔRI) between baseline and treatment intervals for PER2::LUC and [VIP]_e_ rhythms for vehicle and SNAP. For B-E, N=7; G-H, N=6 (WT and KO); J, N=7. In all plots, points represent SCN (paired recordings are joined where relevant), bars and lines/shading are mean±SEM. Scale bar=250μm.

To confirm the specificity of the reporter, VIP-WT and VIP-null SCN were cotransduced with *Syn*.GRAB_VIP1.0_ and *Syn*.jRCaMP1a to enable phasing of [VIP]_e_ against neuronal [Ca^2+^]_i_. Whereas WT SCN displayed robust rhythms in [Ca^2+^]_i_ (Figure 4F), VIP-null SCN displayed damping [Ca^2+^]_i_ rhythms. Furthermore, VIP-null SCN displayed no observable [VIP]_e_ rhythm, with levels remaining at the limit of detection, thereby confirming reporter specificity (Figure 4F). The absence of [VIP]_e_ rhythmicity in VIP-null SCN was reflected in the rhythmicity index (Figure 4G) (repeated-measures two-way ANOVA: Genotype-effect F(1,10)=13.81, p=0.004), where there was a reduced rhythmicity specifically in the [VIP]_e_ report (repeated-measures two-way ANOVA: Reporter-effect F(1,10)=12.52, p=0.005; Interaction F(1,10)=6.70, p=0.03). Consistent with these observations, when [VIP]_e_ was phase-aligned with [Ca^2+^]_i_ and plotted across a 24h interval, VIP-null SCN again lacked a coherent [VIP]_e_ rhythm (Figure 4H), whereas in WT SCN, the [VIP]_e_ peak was directly in phase with neuronal [Ca^2+^]_i_. This is consistent with the phase of peak membrane potential and [Ca^2+^]_i_ in VIP cells (~CT7) (Patton *et al*., 2020) and indicative of daytime neuronal activity driving VIP release. Importantly, this occurs when [GABA]_e_ is at its nadir, consistent with the view that low [GABA]_e_ is permissive for daytime neuronal activity and the dependent neuropeptide release.

To test this, SCN slices expressing GRAB_VIP1.0_ were treated with (S)-SNAP 5114 to examine the effects of loss of GAT3 function on VIP release in the SCN. Following baseline recording, slices treated with vehicle continued to show rhythmic GRAB_VIP1.0_ fluorescence (Figure 4I). In contrast, in slices treated with (S)-SNAP 5114 the GRAB_VIP1.0_ rhythms were immediately compromised resulting in arrhythmicity (Figure 4I), alongside a progressive damping of the PER2::LUC TTFL report. This was reflected in an immediate and severe reduction in the rhythmicity index in the [VIP]_e_ report (repeated-measures two-way ANOVA: Reporter-effect F(1,6)=10.27, p=0.0185) between baseline and treatment intervals (repeated-measures two-way ANOVA: Treatment-effect F(1,6)=16.27, p=0.0069; Interaction F(1,6)=5.62, p=0.0555) (Figure 4J). GAT3 function is therefore necessary to sustain circadian daytime release of the neuropeptide VIP in the SCN.

### Astrocytes initiate network rhythmicity through acute control of [GABA]_e_

Due to the fact that the GAT3 transporter is integral to neuropeptide release from SCN neurons to ensure correct network function, and that it appears to be an astrocyte-enriched transporter and circadian-regulated, we next explored whether the circadian clock of astrocytes could, autonomously, control [GABA]_e_. We have previously used CRY1-complementation targeted to astrocytes to control circadian network function by initiating oscillations in arrhythmic CRY1,2-null SCN (Brancaccio et al., 2019; Patton et al., 2022). Using this approach, however, initiation takes >7 days (Patton *et al*., 2022). To increase the temporal precision with which the astrocytic TTFL could be controlled and its acute effects observed, we utilised translational switching (ts) of CRY1 protein (Maywood et al., 2018). To validate this approach in astrocytes, CRY1,2-null SCN explants with the PER2::LUC reporter were transduced with tsCRY1 (*pCry7*-CRY1(TAG)::mRuby) (Smyllie et al., 2022) alongside the ectopic tRNA-synthetase machinery targeted to astrocytes via *GFAP* promoter (*GFAP*-BFP2-P2A-mMPylS/PylT) (Figure 5A). Following baseline recording, slices were treated with either vehicle or 10mM non-canonical amino-acid alkyne lysine (AlkK), a dose previously shown to initiate CRY1-translation in SCN explants (Maywood *et al*., 2018; McManus et al., 2022; Smyllie *et al*., 2022). Vehicle-treated slices remained arrhythmic. In contrast, TTFL oscillation emerged in 10mM AlkK-treated slices (Figure 5B) with a period (26.95±0.27h) appropriate for CRY1-driven oscillations. It was initiated within 2 days, faster than when Cre-recombinase is used to express CRY1 in astrocytes (Brancaccio *et al*., 2019; Patton *et al*., 2022). In all slices tested, 10mM AlkK initiated SCN-wide rhythms as evidenced by the change in rhythmicity index between baseline and treatment intervals (Figure 5C, ΔRI: vehicle 0.02±0.02 vs 10mM AlkK 0.16±0.04, paired two-tailed t-test t(10)=3.69, p=0.0042). Furthermore, the effect was reversible on withdrawal of AlkK (Supplementary Figure 7).

**Figure 5:**
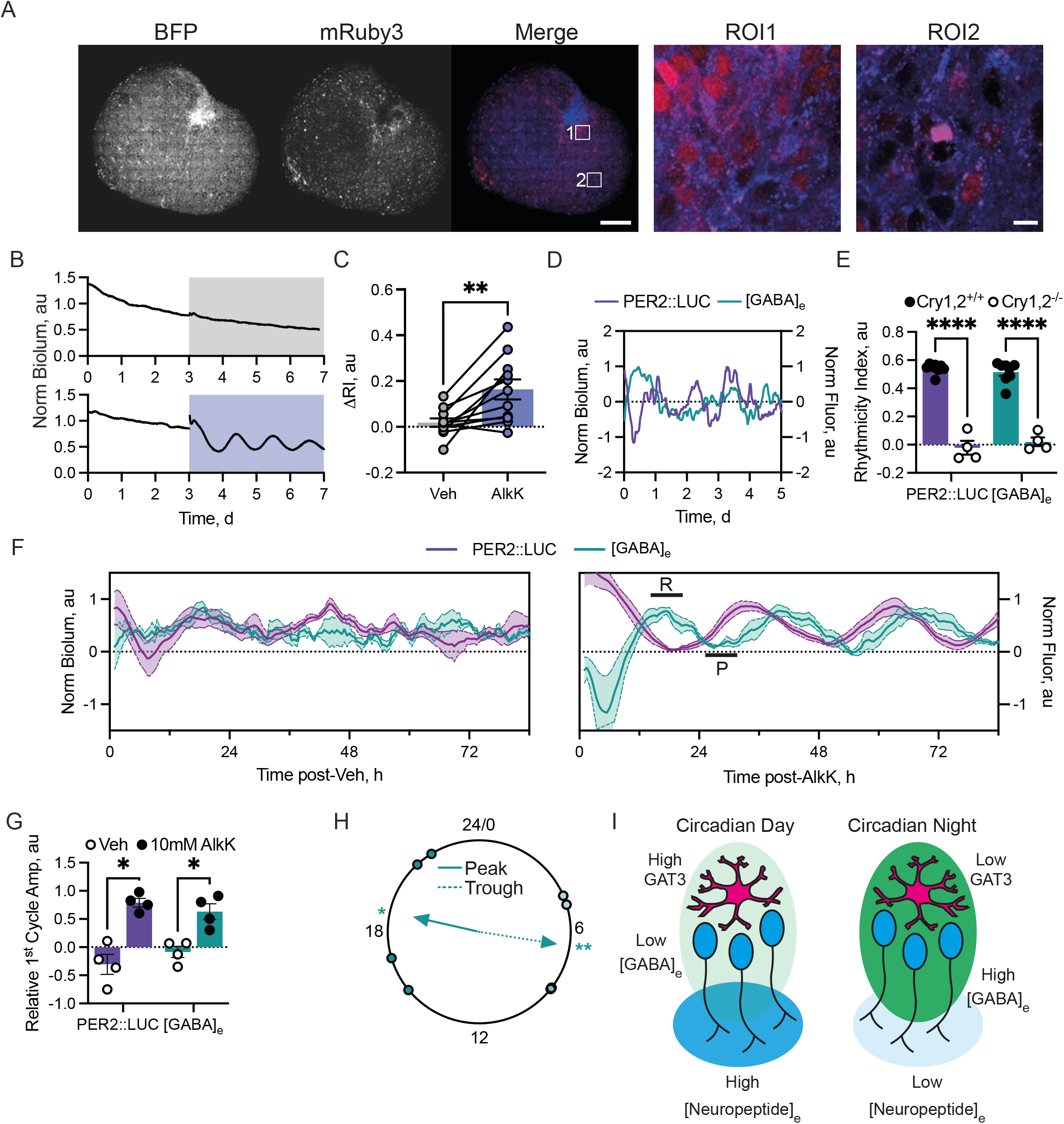
The astrocytic clock acutely controls [GABA]_e_ to transfer circadian information to the SCN network. A. Confocal images of an SCN slice expressing *GFAP*-BFP2-P2A-mMPylS/PylT (left, BFP) and CRY1_TAG_::mRuby3 (middle, mRuby3) following 10mM AlkK treatment alongside a false-coloured merged image (right, Merge, BFP (blue) and mRuby3 (red)). Two zoomed-in ROIs are shown to the right. B. PER2::LUC traces from vehicle (upper, grey) or 10mM AlkK-treated (lower, purple) CRY1,2-null SCN. C. Change in rhythmicity index (ΔRI) between baseline and treatment intervals vehicle- or 10mM AlkK-treated SCN. D. Normalised PER2::LUC and [GABA]_e_ emissions from a CRY1,2-null SCN. E. Histogram showing rhythmicity index measures for PER2::LUC and [GABA]_e_ from wild-type (filled circles, CRY1,2^+/+^, data from Figure 1C) and CRY1,2-null (open circles, CRY1^-/-^,2^-/-^) SCN. F. PER2::LUC and [GABA]_e_ traces showing their dynamics during vehicle (left) or AlkK-treatment (right). In the right-hand plot, phases where GABA is repressive or permissive are indicated by bars labelled R or P, respectively. G. Histogram showing relative amplitude of the first cycle following treatment for PER2::LUC and [GABA]_e_ with vehicle (open circles) or 10mM AlkK (filled circles). H. Rayleigh plot showing timing of the first cycle trough (dotted line) and peak (solid line). I. Schematic describing how the astrocytic clock controls network function via [GABA]_e_ flux. During circadian day (left), astrocytic GAT3 expression is high, resulting in lower [GABA]_e_ (light green shading). This leads to increased neuronal activity and neuropeptide release (blue shading). During circadian night (right), astrocytic GAT3 expression is low, resulting in high [GABA]_e_ (green shading) leading to reduced neuronal activity and neuropeptide release (light blue shading). In all plots, points represent SCN with paired measures joined where relevant. Bars and lines/shading represent mean±SEM. For C N=11, E N=8 Wild-type and 4 CRY1,2-null, G/H N=4. Scale bar=200μm/10μm.

This approach therefore presents a unique opportunity to observe acute [GABA]_e_ changes as network oscillations are initiated by the cell-autonomous astrocytic TTFL. CRY1,2-null slices were super-transduced with *Syn*.iGABASnFR to monitor [GABA]_e_ as network rhythms initiated. In these CRY1,2-null SCN slices, similar to the TTFL rhythm, [GABA]_e_ was arrhythmic (Figure 5D, E) (repeated-measures two-way ANOVA: Reporter-effect F(1,10)=0.11, p=0.75) with a greatly reduced rhythmicity index in CRY1,2-null versus wild-type slices (Figure 1C, Figure 5E, repeated-measures twoway ANOVA: Genotype-effect F(1,10)=300, p<0.0001; Interaction: F(1,10)=1.61, p=0.23). Arrhythmicity was maintained following vehicle-treatment (Figure 5F). In contrast, under 10mM AlkK-treatment, not only were bioluminescent rhythms initiated, but [GABA]_e_ also started to fluctuate with an initial rapid decrease followed by a rapid increase into a sustained peak (Figure 5F, marked R). This sustained elevation of [GABA]_e_ aligned with the nadir of the bioluminescent signal, marking the first circadian night and repressive phase of the TTFL clock (Figure 5F). [GABA]_e_ then fell into a sharp trough (Figure 5F, marked P) in the first circadian day, marked by rising PER2::LUC bioluminescence. Subsequently, [GABA]_e_ rose again to another sustained peak in the second circadian night (>CT12). These dynamics, therefore, align low [GABA]_e_ with the positive arm of the neuronal TTFL (Brancaccio *et al*., 2019), and high [GABA]_e_ with the repressive arm. As network time-keeping is initiated, acute changes in [GABA]_e_ driven by the astrocytic TTFL generate network oscillation. They first facilitate repression via neuronal inhibition under high [GABA]_e_, which is then followed by co-ordinated facilitation of neuronal activity under low [GABA]_e_ via the permissive uptake of GABA from the extracellular space.

To assess the initiation of TTFL and [GABA]_e_ dynamics quantitively, we measured the coherence of the initiated oscillations by calculating their relative amplitudes, defined as the difference between the peaks and troughs predicted from the dynamics of the PER2::LUC bioluminescence recorded in PMTs (Figure 5B) and the circadian dynamics of [GABA]_e_ in wild-type slices (Figure 1). We saw that in the high-temporal resolution PMT traces, PER2::LUC reached its first initiated peak 34.5h (34.6±0.6h, Figure 6B) after AlkK-treatment. Using this, we identified when the troughs in the PER2::LUC and [GABA]_e_ oscillation that preceded this peak would have fallen (21h and 26h post-treatment, respectively) and where the ensuing peak in [GABA]_e_ would have fallen (39.5h post-treatment) in the normalised data (Figure 5F). Using these measures, we observed a low relative amplitude of the vehicle-treated oscillation in PER2::LUC and [GABA]_e_, which increased in both when treated with AlkK (Figure 5G) (repeated-measures two-way ANOVA: Reporter-effect F(1,3)=0.04, p=0.85; Treatment-effect F(1,3)=36.58, p=0.009; Interaction F(1,3)=0.18, p=0.18). This indicates that the peak and trough of the [GABA]_e_ oscillation driven by the astrocytic clock were initiated immediately with dynamics corresponding directly with the [GABA]_e_ oscillation in a competent SCN (Figure 1A, Figure 5H) (Rayleigh test against specified mean direction (μ): peak (μ=CT19.7) p=0.017, R=0.73; trough (μ=CT8.2) p=0.009, R=0.80), and importantly occur acutely as network oscillations initiated. Thus, [GABA]_e_ is controlled by the cell-autonomous TTFL of astrocytes and encodes and transfers circadian information directly to the SCN neuronal network.

## Discussion

Free-running SCN explants display robust rhythms in [GABA]_e_, peaking at night in antiphase to neuronal activity (Figure 1). This observation is paradoxical because a peak in [GABA]_e_ would be expected in circadian day when SCN GABAergic neurons are active. Equally, GABA is inhibitory, and so high daytime [GABA]_e_ would be expected to suppress neuronal activity and therefore its own release. The paradox could be resolved if GABA were excitatory, a condition observed during development (Ben-Ari et al., 2012), arising when intracellular chloride levels ([Cl^+^]_i_) are high. This is achieved via changes in the expression of K-Cl and Na-K-2Cl co-transporters and leads to a reversal of GABA_A_-receptors on activation (Kaila et al., 2014; Payne et al., 2003). Indeed, a model whereby SCN neurons may switch GABAergic transmission between excitatory and inhibitory on a circadian basis has been proposed (DeWoskin et al., 2015) and excitatory GABAergic drive reported in the SCN under certain conditions. There is, however, no consensus on its temporal or spatial patterning (for review see (Albers *et al*., 2017)). More directly, our data contradict this model, indicating that GABAergic signalling, mediated by GABA_A_-receptors, forms a suppressive rather than excitatory axis (Figure 2, Supplementary Figure 2). This suppression is presumably driven by extra-synaptic GABA_A_-receptors which are well-suited to sense circadian changes in bulk [GABA]_e_ and drive long-term inhibition during the night due to their high affinity and decreased desensitisation in the prolonged presence of GABA (Albers *et al*., 2017; Brickley and Mody, 2012; Farrant and Nusser, 2005).

Our observations, provide an alternative resolution of the paradox. In our model astrocytes, not neurons, control [GABA]_e_ (Figure 5I). This predicts that GABA uptake during the day (when neurons are most actively releasing GABA) via the GAT3 transporter prevents GABA spill-over from synaptic sites and accounts for the sharp trough in the [GABA]_e_ oscillation (Figure 1). This uptake is permissive to neuronal activity, which in turn supports electrically sensitive CREs in *Per* genes, thereby leading to robust high amplitude TTFL oscillation across the SCN (Figure 2). Conversely, elevated [GABA]_e_ sustains the nocturnal repressive phase of the TTFL. The critical daytime uptake of GABA was confirmed by direct imaging of the [GABA]_e_ oscillation under treatment with the GAT3-specific inhibitor (S)-SNAP 5114, where loss of rhythmicity in the GABA report is caused by an acute elevation of the trough without effects on peak level (Figure 3). Day-time removal of GABA from the extracellular milieu is necessary for neuronal activity, and activity-dependent neuropeptide release, which reinforces circuit-level rhythms (Figure 4).

How then, does the [GABA]_e_ rhythm sit in antiphase to the activity of its presumed source? Using our published single-cell transcriptomic dataset (Morris *et al*., 2021) we found a day-night difference in GAT expression, predominantly on astrocytes (Figure 3, Supplementary Figure 3). SCN expression of GATs has been reported previously (Barca-Mayo *et al*., 2017; Coomans *et al*., 2021; Moldavan *et al*., 2015) and potential circadian variation shown in rat SCN (Moldavan *et al*., 2015) and cortical astrocytes (Barca-Mayo *et al*., 2017). Our functional tests of GAT1 and GAT3 inhibition revealed that their inactivation suppressed the PER2::LUC oscillation (Figure 3), because of the consequent rising levels of GABA_A_-receptor activation (Figure 2). Furthermore, GAT3 inhibition produced stronger effects on application and during washout (Figure 3, Supplementary Figure 4), likely due to the fact that GAT3 has a higher affinity for GABA compared to GAT1 (Zhou and Danbolt, 2013) allowing GAT3 to more efficiently lower GABA levels in the absence of GAT1 function. We therefore consider GAT3 to be the principal transporter for controlling [GABA]_e_ in the SCN and thereby influencing SCN function.

Consistent with our model, whereby the daytime clearance of [GABA]_e_ is permissive to time-keeping, sustained loss of GABAergic release from neurons by genetic targeting of the vesicular transporter VGAT does not markedly influence SCN molecular rhythmicity (Ono *et al*., 2019), even though it compromises transmission of behaviourally relevant cues at distal extra-SCN targets. Equally, TTFL function in the SCN is sustained during blockade of GABA_A_- and/or GABA_B_-receptors (Figure 2, (Freeman *et al*., 2013b; Patton *et al*., 2016)), although GABA_A_ blockade ((+)-bicuculline) elevates both the peak and trough of the oscillation (Figure 2), indicating that GABA fine-tunes SCN electrical activity within a certain dynamic range that complements TTFL function (Maejima *et al*., 2021; Ono *et al*., 2019). The nocturnal rise in [GABA]_e_ is important because it curtails the permissive low-GABA state. If that low-GABA state is prevented by GAT3 blockade, not only is the neuronal activity rhythm (monitored by [Ca^2+^]_i_) damped, but the consequent release of neuropeptides is curtailed.

VIP is a principal SCN neuropeptide, essential for circadian neuronal synchrony and high-precision oscillation (Colwell et al., 2003; Harmar et al., 2002; Maywood et al., 2006), driving the cell-autonomous TTFL of cells expressing its receptor, VPAC2, by kinase-dependent signalling to CRE-elements in the *Per* genes (Hamnett et al., 2019; Hastings *et al*., 2018). A VIP-GABA counterbalance mechanism has been proposed (Barca-Mayo *et al*., 2017; Freeman *et al*., 2013a), and GAT-mediated GABA uptake suggested as a conduit for circadian astrocyte-to-neuron information transfer in coculture *in vitro* models (Barca-Mayo *et al*., 2017). Indeed, pharmacological disruption of GABAergic signalling counteracts some of the deficits of VIP deficiency in isolated SCN (Freeman *et al*., 2013a), adding to the notion that opposing GABAergic and neuropeptidergic cues set the dynamic range for normal TTFL function. Using a VIP-specific reporter (Wang *et al*., 2022), we mapped peak VIP release to the middle of circadian day (CT6.97), consistent with previous recordings and the known activity cycle of VIP neurons (Ono *et al*., 2023; Patton *et al*., 2020), and showed that VIP and GABA are ordinarily maintained in a distinct temporal segregation in the SCN, held in antiphase by GABA uptake by astrocytic transporters. Compromising this uptake rapidly damped rhythmic VIP release, revealing explicitly the interplay between [GABA]_e_ and [VIP]_e_ in circadian time, and showing that control of [GABA]_e_ not only directs the cell-autonomous TTFL, but also circuit-level SCN time-keeping.

Night-time release of GABA is compatible with our previously proposed astrocyte-to-neuron signalling axis whereby nocturnally-released astrocytic glutamate depolarises neurons expressing NR2C-containing NMDA receptors, increasing action potential-independent GABA release from these neurons (Brancaccio *et al*., 2017). The effectiveness of this synaptic release would be augmented by inactivation of GABA uptake, allowing for enhanced spill-over of GABA into extra-synaptic sites. In these complementary, co-ordinated ways, astrocytes can suppress the SCN neuronal network during the night directly by manipulating the properties of their extracellular milieu.

Evidence of this reciprocal astrocyte-to-neuron signalling is provided by the re-emergence of TTFL rhythms following the removal of GAT3 inhibition: on washout the TTFL amplitude did not return immediately. Rather, it grew steadily, cycle-to-cycle (Supplementary Figure 4B, C). This progressive recovery following a switch from high [GABA]_e_ to low [GABA]_e_ can be explained by the mutual dependence of neuronal and astrocytic TTFLs. In the prolonged high [GABA]_e_ state, loss of neuronal rhythmicity likely also impairs astrocytic rhythms, and following transfer to the low [GABA]_e_ state, it requires several cycles of mutual reinforcement to establish full spectrum oscillations in both cell-types. A comparable re-establishment of TTFL cycles occurs following washout of cycloheximide (suspending cell-autonomous TTFLs) or tetrodotoxin (suspending neuronal signalling) (Abel et al., 2016; Yamaguchi *et al*., 2003; Yamaguchi et al., 2013). In the case of GAT3 inhibition, these dynamics are not, however, driven purely by release of GABAergic inhibition to the neurons: washout of muscimol does not produce the same continuous increase in amplitude (Supplementary Figure 1). The building of amplitude upon washout of GAT3 therefore represents a gradual reconfiguration of the mechanisms to control GABA uptake and release across neurons and astrocytes following chronic inactivation of an astrocyte-specific transporter. We therefore interpret this as the SCN network iteratively rebalancing itself as the neurons and astrocytes are re-coupled on a cycle-to-cycle basis. Comparable washout dynamics are seen following disruption of astrocyte-to-neuron signalling when NR2C-receptors or glutamate uptake are inhibited and then restored (Brancaccio et al 2017).

Our model proposes that daily rhythms in GABA uptake, driven principally by astrocytes, control circadian dynamics of bulk GABA in the extracellular space. This uptake is dependent on a functional SCN clock, as [GABA]_e_ is arrhythmic in CRY1,2-null explants, consistent with previous observations of arrhythmic astrocytic TTFL and [Ca^2+^]_i_, and extracellular glutamate (Brancaccio *et al*., 2019; Marpegan et al., 2011; Patton *et al*., 2022). In order to identify causal events in a repetitive, cyclical system such as the SCN, where changes on one cycle may be caused by events some cycles previously, it is necessary to have precise temporal control of the pertinent components. We therefore used tsCRY1-expression specifically in astrocytes to activate their cell-autonomous TTFL and thereby test our model by following the sequence of events (Figure 5) as circadian rhythmicity was initiated, *de novo*, in arrhythmic CRY1,2-null SCN (Maywood *et al*., 2018; McManus *et al*., 2022; Smyllie *et al*., 2022). This revealed precise patterning of coherent [GABA]_e_ dynamics occurring over the first 48h, with changes in [GABA]_e_ preceding those in the TTFL. Furthermore, from their initiation stage, these rhythms matched the relative phasing in wild-type SCN. Thus, in the absence of a functional clock in the rest of the SCN, astrocytes can impose their cell-autonomous circadian state onto the SCN network by initiating circadian cycles of [GABA]_e_. We have, therefore, revealed a new level of astrocyte-to-neuron communication that controls SCN network dynamics on a circadian timescale, definitively showing that astrocytes augment neuronal function. Having established this mode of communication, it is now imperative to explore whether further signalling axes support this and to determine the mechanisms by which neurons reciprocate this signalling to astrocytes. A particularly striking observation from our results is that despite neurons being the (presumed) source of GABA in the SCN network, circadian control of this signalling is devolved to the astrocytes. Is this, therefore, a more general mechanism whereby astrocytes regulate other GABAergic circuits in the brain across different time-scales?

## Acknowledgements

The authors would like to thank J.E. Chesham, O. Johnson, LMB Biomed and Ares facilities, and Genotyping staff for animal husbandry support. The authors also thank Mechanical and Electronics Workshops and Light Microscopy Facility at the MRC Laboratory of Molecular Biology for technical support. Supported by core funding from Medical Research Council (MRC), as part of United Kingdom Research and Innovation (also known as UK Research and Innovation) (MRC File Reference No. MC_U105170643) to M.H.H and BBSRC Project Grant (BB/R016658/1) to M.H.H and A.P.P. For the purpose of open access, the author has applied a CC BY public copyright licence to any Author Accepted Manuscript version arising.

## Author Contributions

A.P.P and M.H.H designed the study and wrote the manuscript. A.P.P performed the experiments and analysed the data. E.L.M. provided scRNA-seq data. D.M, J.W.C, Y.L and H.W. provided novel reagents.

## Methods

### Resource availability

Further information and requests for reagents should be directed to and will be fulfilled by the lead contact, Michael Hastings.

### Materials availability

Plasmids generated in this study are available upon request as detailed above.

### Data and code availability

The publicly available single cell RNA-seq data were accessed from GEO under accession number GSE167927 (Morris *et al*., 2021). The R script used to analyse the single-cell data is included in the supplementary information. Any additional data or information is available from the lead contact upon request.

### Experimental model and subject details

#### Animals

All experiments were performed in accordance with the UK animals (Scientific Procedures) Act of 1986, with local ethical approval (MRC LMB AWERB). PER2::Luciferase (PER2::LUC) mice were kindly supplied by J.S. Takahashi University of Texas Southwestern Medical Center, Dallas, TX; (Yoo et al., 2004). VIP-null mice were a gift from C.S. Colwell; (Colwell *et al*., 2003). CRY1,2-null mice were derived from founders supplied by G. van der Horst (Erasmus University Medical Center, Rotterdam, The Netherlands; (van der Horst et al., 1999)). CRY1,2-null mice were crossed to the PER2::LUC line in-house and all mice were maintained on a C57BL6/J background.

#### Organotypic slice preparation and AAV transduction

Postnatal day 10 (P10) to P12 mice of either sex were killed according to local and UK Home Office approved rules. The brains were removed and transferred to ice-cold GBSS supplemented with 5mg/ml glucose, 3mM MgCl2, 0.05mM DL-AP5 and 100nM (+)-MK-801. Brains were trimmed into a block containing the hypothalamus before 300μm thick coronal slices were made on a tissue chopper (McIlwain, UK). The SCN was then dissected free from the slice and cultured as an organotypic explant via the interface method. Slices were rested in culture medium supplemented with 3mM MgCl2, 0.05mM before being transferred to medium for 1 week before use (Hastings et al., 2005).

### Method details

#### AAVs and molecular biology

The following AAVs were obtained as viral preps directly from Addgene. pAAV.*hSynap*.iGABASnFR (*Syn*-iGABASnFR, AAV1) was a gift from Loren Looger (Addgene viral prep # 112159-AAV1; http://n2t.net/addgene:112159; RRID:Addgene_112159). pAAV.*Syn*.NES.jRCaMP1a.WPRE.SV40 (*Syn-*jRCaMP1a, AAV1) was a gift from Douglas Kim & GENIE Project (Addgene viral prep # 100848-AAV1; http://n2t.net/addgene:100848; RRID:Addgene_100848).

*hSyn*.GRAB_VIP1.0_.WPRE was a gift from Yulong Li (School of Life Sciences, Peking University, China) and was obtained as a plasmid before being packaged into AAV serotype 9 viral particles by VectorBuilder (https://www.vectorbuilder.com/). tsCRY1::mRuby (*pCry1*-CRY1(TAG)::mRuby3) was produced in-house and packaged by VectorBuilder as AAV1 serotype AAV particles as detailed previously (Smyllie *et al*., 2022). *GFAP*-BFP2-P2A-MmPylS was made in-house by swapping the fluorescent tag in the *GFAP*-mCherry-P2A-MmPylS plasmid (Krogager et al., 2018) for BFP2 before being packaged by VectorBuilder as AAV serotype 5 viral particles.

#### AAV transduction

In the case of SCN slices that received AAV transductions, following at least a week in culture, the medium was changed and slices were transduced with 1-2μl of AAV (>1×10^12^ genome copies/ml in PBS) applied directly as a droplet to the top of the SCN. In the case of SCN receiving multiple serial transductions, 2-3 days were left between each round of transduction. A week following the final transduction, the medium was changed.

#### Real-time bioluminescent and fluorescent imaging

Aggregate bioluminescence was monitored in real-time in customised light-tight incubators fitted with individual photon-multiplier tubes (PMTs; catalog #H9319-11 photon counting head, Hamamatsu). Data were binned into 6-minute epochs for export and analysis. Slices were maintained in a HEPES-buffered medium containing DMEM, supplemented with Glutamax, penicillin/streptomycin, FCS, B27 and 100μM luciferin as detailed previously (Hastings *et al*., 2005). Dishes were sealed with glass coverslips.

For combined circadian fluorescent/bioluminescent imaging, SCN slices were sealed in glass-bottomed imaging dishes (P35G-0-10-C, Mattek) containing the same medium as in PMT recordings. Slices were imaged on an LV200 Bioluminescence Imaging System (Olympus). Bioluminescence was acquired for between 9.5 and 29.5 mins, while multiplexed fluorescence acquisition was set at 100ms. The acquisition intervals for combined circadian imaging were 30 mins.

#### Immunostaining and Confocal Microscopy

Membrane-attached SCN slices were excised via scalpel and fixed by submersion in 4% paraformaldehyde (PFA) diluted in phosphate buffer for 30 minutes at room temperature. Following this, slices were washed twice for 10 minutes each in 0.01M phosphate buffered saline (PBS) at room temperature. Washed slices were transferred to a blocking buffer composed of 0.01M PBS supplemented with 1% bovine serum albumin (BSA), 0.5% Triton X-100 and 5% Normal Goat Serum (Day 1 buffer) and blocked for 6 hours at room temperature before being transferred to Day 1 buffer supplemented with primary antibodies/antisera and incubated overnight at 4°C. Following incubation with the primary antibodies/antisera, slices were washed twice in Day 1 buffer diluted 1/3 in 0.01M PBS (Day 2 buffer) for 15 minutes each, at room temperature. Slices were then transferred to the Day 2 buffer supplemented with the secondary antibodies and incubated for 1 hour at room temperature. Following incubation with the secondary antibodies, slices were washed twice in Day 2 buffer and twice in 0.01M PBS for 15 minutes each at room temperature. Finally, slices were rinsed in ddH2O and mounted on glass slides before being coverslipped with Vectamount HardSet antifade mounting medium (H-1400, Vector Laboratories). Where fluorophores permitted, the mounting medium used was supplemented with DAPI (H-1500, Vector Laboratories). All incubation and wash steps were carried out on a shaker.

The following primary antisera were used at the following dilutions: Rabbit anti-GAT1 (Synaptic Systems, 274 102) (1:500) and Guinea Pig anti-GAT3 (Synaptic Systems, 274 304) (1:500). The following secondary antibodies were used at the following dilutions: Goat anti-Rabbit Alexa 488 (Invitrogen, A11008) (1:1000) and Goat anti-Guinea Pig 568 (Invitrogen, A11075) (1:1000).

Mounted immunostained SCN were imaged on a Zeiss 880 Airyscan confocal microscope controlled by Zen software (Zen 2.3, Zeiss). Imaging was carried out with a 20x apochromatic objective and the whole of the SCN slice was captured in tiles at 3 different focal planes due to the uneven thickness of cultured SCN explants. Post-imaging, Z-projections were made of the maximum pixel intensity to create a composite image of the entire SCN explant.

In the case of imaging native fluorescent markers expressed via AAVs, SCN were fixed, washed and mounted before being imaged on a Zeiss 880 Airyscan confocal microscope with a 63x oil-immersion apochromatic objective to enable better resolution imaging of the fluorescent proteins. In all cases, slices were imaged as tiles to capture the whole of the SCN.

#### Pharmacological treatments

(R)-baclofen, (+)-bicuculline, CI 966 HCl, DMSO, muscimol, SCH 50911 and (S)-SNAP 5114 were obtained from Tocris, UK. (R)-baclofen and muscimol were solubilised in water to stock solutions of 20mM and 100mM respectively. (+)-bicuculline, CI 966 HCl, SCH 50911 and (S)-SNAP 5114 were solubilised in DMSO at a stock concentration of 100mM (with the exception of CI 966 HCl which was at 50mM). The corresponding vehicle (water or DMSO) was used with the corresponding pharmacological agent and for dose-response curves, dilutions were made in the same vehicle as serial dilutions. Pharmacological agents were washed out by transferring the membrane and SCN to fresh media three times serially, waiting for 10 minutes at each step.

Alkyne lysine (AlkK, N6-2-propynyloxycarbonyl-l-lysine, synthesised in-house) was prepared freshly as a stock solution of 100mM dissolved directly in recording medium, before being adjusted to pH7.0 by addition of NaOH, as previously described (Maywood *et al*., 2018; McManus *et al*., 2022; Smyllie *et al*., 2022). The AlkK solution was then sterile filtered before being added to slice medium at a 1:10 dilution to result in a final concentration of 10mM in the recording medium. For vehicle treatments, a 1:10 dilution of medium without AlkK was added to the recording medium. AlkK or vehicle were washed out of the slices by transferring the membrane and SCN to fresh media six times serially, waiting for 10 minutes at each step.

### Quantification and statistical analysis

#### Analysis of real-time bioluminescent and fluorescent imaging

For PMT data, peaks and troughs were identified from the raw bioluminescence in wild-type oscillations. For CRY1,2-null experiments, PMT data was detrended by subtracting a polynomial fit before an FFT-NLLS fit was made in BioDARE2 (Zielinski et al., 2014) (https://biodare2.ed.ac.uk/) in order to assess circadian properties. Rhythmicity index was calculated by taking the mean of the autocorrelation of the detrended time series at 26, 52 and 78h, a cycle length appropriate for CRY1-driven oscillations. The autocorrelation was calculated in R using the acf command in the base R stats package.

All manipulation of real-time bioluminescence and fluorescence images was carried out in FIJI (Schindelin et al., 2012). Aggregate bioluminescence and fluorescence were analysed by exporting sequential images to TIFFs. Bioluminescence stacks were then de-noised by removing outliers and the raw mean grey values through the stacks were exported. For fluorescence, stacks were background subtracted by applying the built-in background subtraction command, using the rolling ball algorithm and setting the size to 5 pixels before the raw mean grey values were exported. Exported data was further analysed in Excel and R and detrended by subtracting a polynomial fit in order to identify peaks and troughs and rhythmicity indexes. Under some circumstances, peak phases, periods and relative amplitude error was identified using FFT-NLLS measures in BioDARE2 as above. Rhythmicity index was calculated as above, using the autocorrelation at 24, 48 or 72h for wild-type oscillations.

Rayleigh statistics were calculated using the r.test command in the CircStats package in R (version 0.2-6). To compare the phases of the astrocyte-initiated [GABA]_e_ peak and trough with the wild-type phases, a modified version of the Rayleigh test was used where the reported phase was tested against an alternative hypothesis where the wild-type peak and trough timings were supplied as specified mean directions (in radians) using the v0.test command in the CircStats package.

For GRAB_VIP1.0_ recordings in VIP-WT and VIP-KO SCN, jRCaMP1a and GRAB_VIP1.0_ data were detrended before being normalised and overlaid as 24h intervals centred around the peak of the jRCaMP1a signal.

In order to assess the acute changes in [GABA]_e_ dynamics in response to GAT3 inhibitor treatment, the peaks and troughs were projected forward by fitting a line to the pre-treatment peaks or troughs for vehicle and inhibitor treatment in the raw aggregate trace. This allowed prediction of the level at which these parameters would have been to allow comparison with the recorded values, which was expressed as a ratio of the actual amplitude/predicted amplitude.

#### Analysis of previously published scRNA-seq datasets

Data from (Morris *et al*., 2021) were accessed from NCBI Gene Expression Omnibus with the accession number: GSE167927. Data were analysed in R using the Seurat package (Seurat version 4.0.5, (Hao et al., 2021)). A script of the analysis is available in supplementary information.

#### Experimental design and statistical analysis

Where possible, slices received paired treatments (vehicle and drug) and were exposed to all concentrations of drug along a dose-response curve. Where this was not possible, or slices died during the course of the experiment, slices were assigned randomly to groups. All data were analyzed in Excel (Microsoft), R (version 3.6.1; R Foundation for Statistical Computing), RStudio (version 1.2.1335, RStudio Inc.) and GraphPad Prism 9 (GraphPad). All the statistical tests used are listed in the text and figure legends. All numbers reported in text are mean±SEM unless otherwise stated.

