## Supplementary Materials for "Astrocytic control of extra-cellular GABA drives circadian time-keeping in the suprachiasmatic nucleus"

**Supplementary Figure 1: iGABASnFR fluorescence is membrane targeted across the SCN**

Confocal images of an SCN slice expressing neuronal-targeted iGABASnFR (left, iGABASnFR) alongside nuclear DAPI staining (middle, DAPI) and a false-coloured merged image (right, Merge, iGABASnFR (green) and DAPI (blue)). Two zoomed-in ROIs are shown to the right. Scale bar=200μm/10μm.

Supplementary Figure 1

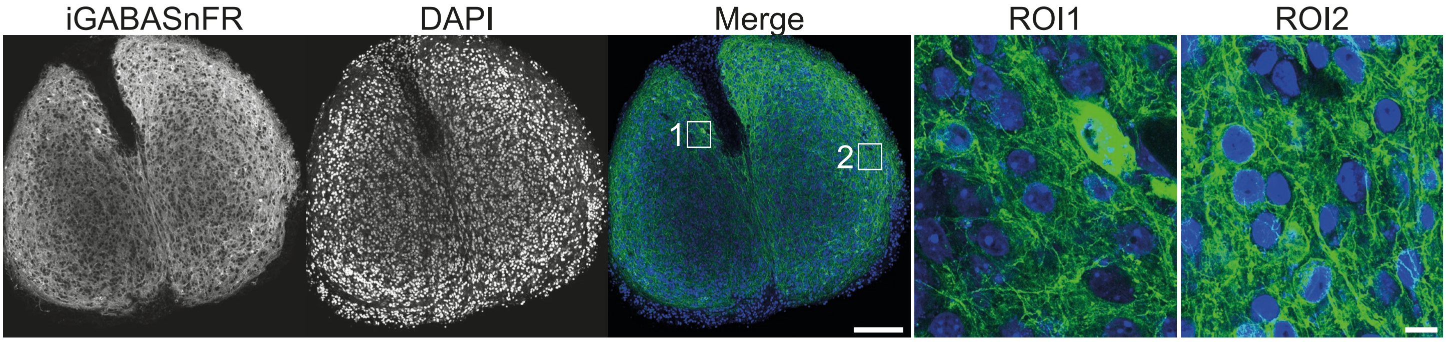

**Supplementary Figure 2: The effects of chronic muscimol and (+)-bicuculline treatment are reversible on washout**

A. Example normalised PMT traces showing PER2::LUC bioluminescence before, during and after treatment with 100μM muscimol (yellow). B. Example normalised PMT traces showing PER2::LUC bioluminescence before, during and after treatment with 100μM (+)-bicuculline (purple).

Supplementary Figure 2

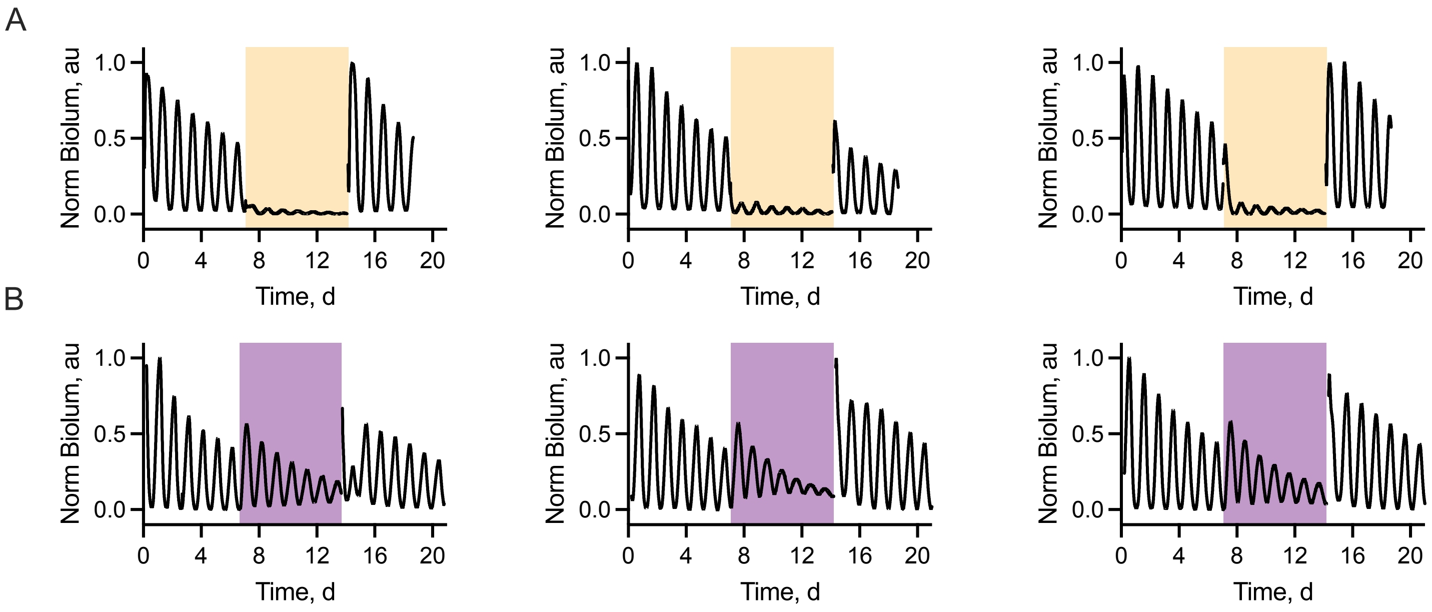

**Supplementary Figure 3: Effects of chronic treatment of SCN slices with $GABA_A$ or $GABA_B$ receptor agonists and antagonists on peak and trough levels**

Histograms showing PER2::LUC trough (left) or peak (right) levels of the treatment interval normalised to the baseline interval for slices treated with different concentrations of (A) muscimol, (B) (R)-baclofen, (C) (+)-bicuculline, and (D) SCH5011. In all plots, individual points represent single SCN and bars are mean±SEM.

Supplementary Figure 3

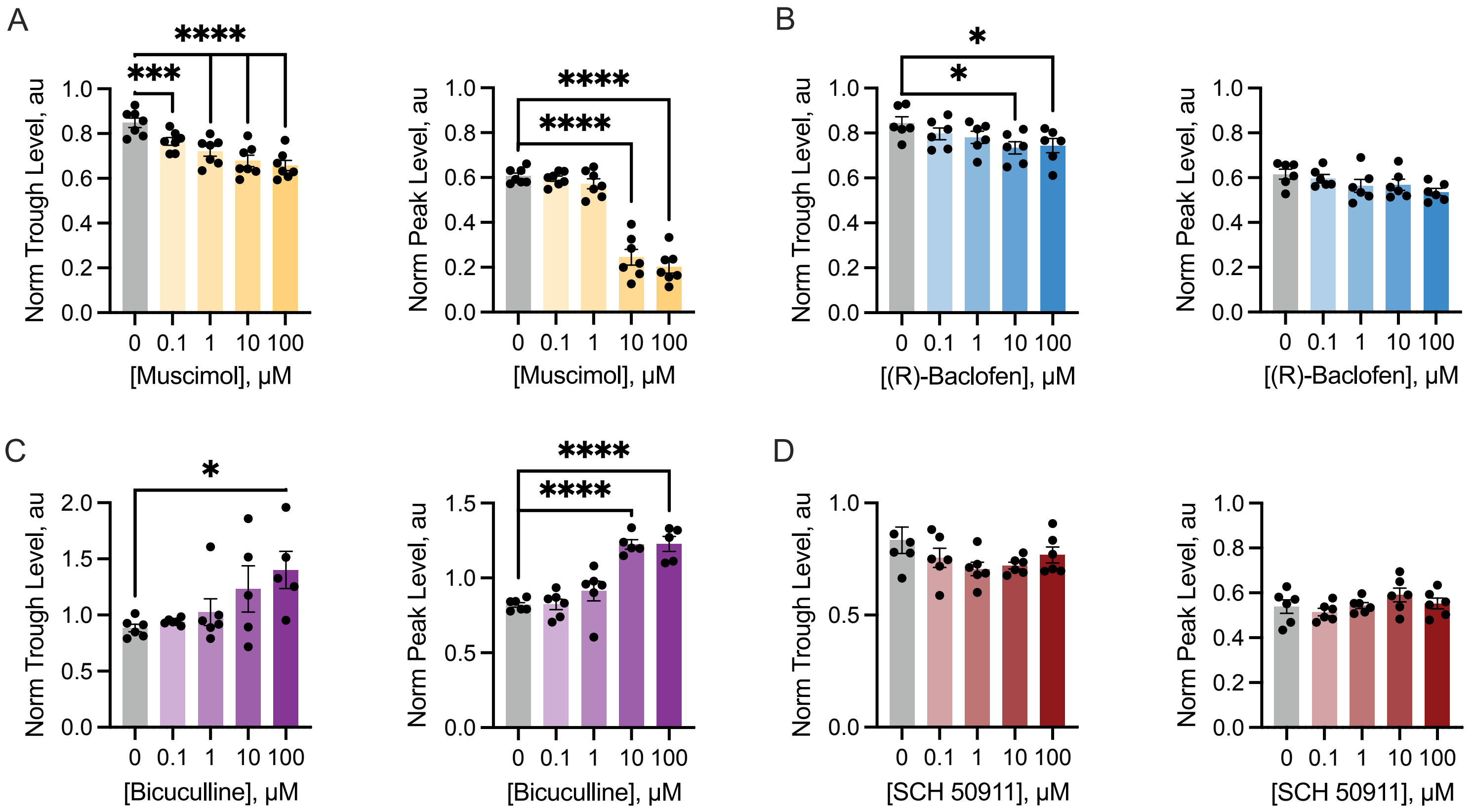

**Supplementary Figure 4: GAT1 and GAT3 expression within SCN explants**

Confocal micrographs showing SCN explants immunostained for the GABA transporters GAT1 (left) and GAT3 (middle) alongside the nuclear stain DAPI (right). Scale bar=200µm. B. Histograms showing the relative expressions of GAT1 (*Slc6a1*, left) and GAT3 (*Slc6a11*, right) relative to the house-keeping gene *Rn18s* in slices harvested during circadian day (CT0-4) (Day, orange) or circadian night (CT12-16) (Night, blue). N=4 day/7 night.

Supplementary Figure 4

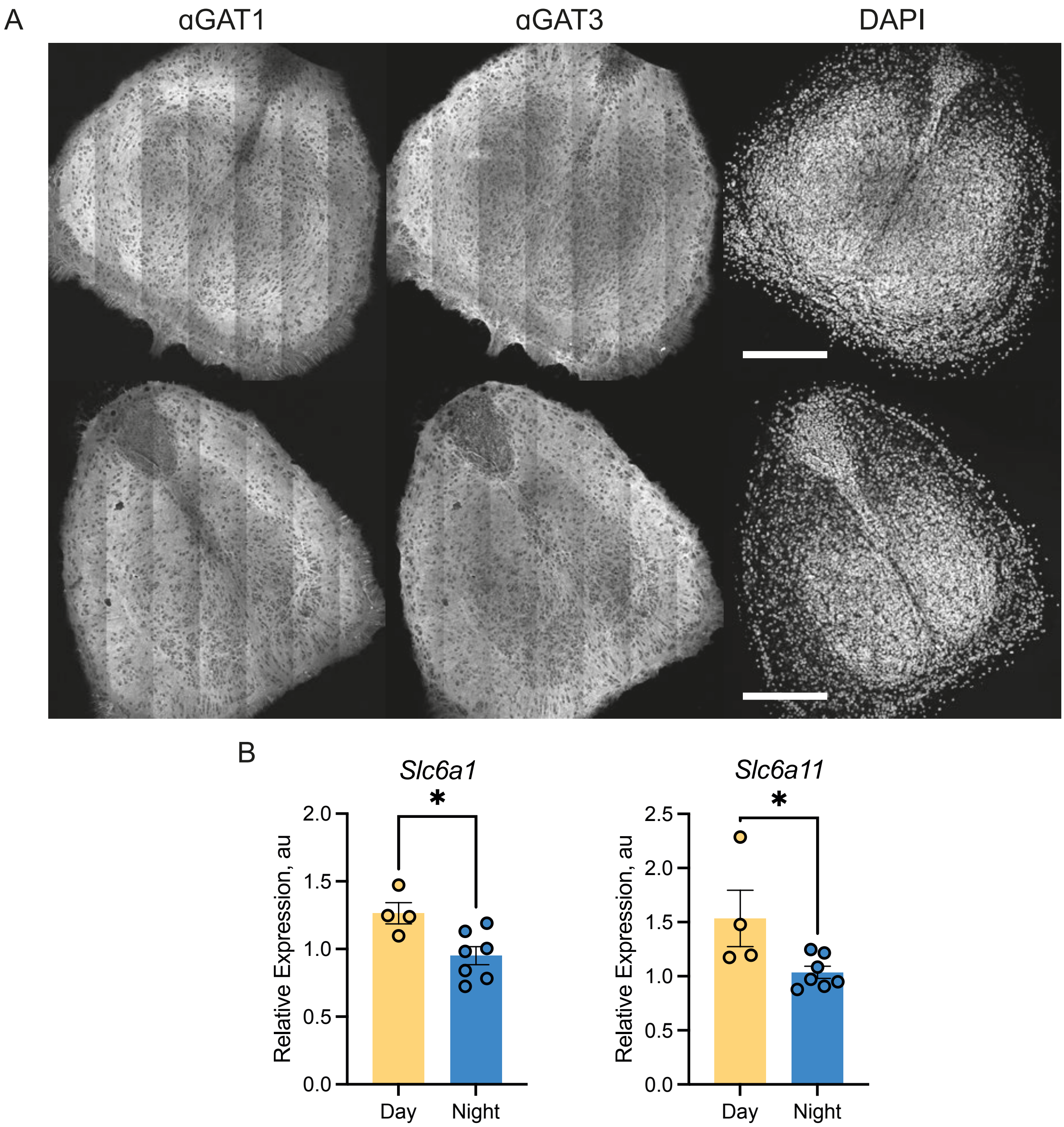

**Supplementary Figure 5: The effects of CI 966 HCl and (S)-SNAP 5114 treatment are reversible upon washout**

A. Example normalised PMT traces showing PER2::LUC bioluminescence before, during and after treatment with 50μM CI-966 HCl (green). B. Example normalised PMT traces showing PER2::LUC bioluminescence before, during and after treatment with 50μM (S)-SNAP 5114 (orange). C. Normalised cycle-to-cycle amplitude of the PER2::LUC oscillation post-washout for slices treated with DMSO vehicle or 50μM (S)-SNAP 5114. N=6 vehicle/4 (S)-SNAP 5114. Two-way ANOVA: Time effect: $F(4,31)=5.74$, $p=0.001$; Treatment effect: $F(1,8)=8.85$, $p=0.018$; Time x Treatment interaction: $F(4,31)=8.72$, $p<0.0001$.

Supplementary Figure 5

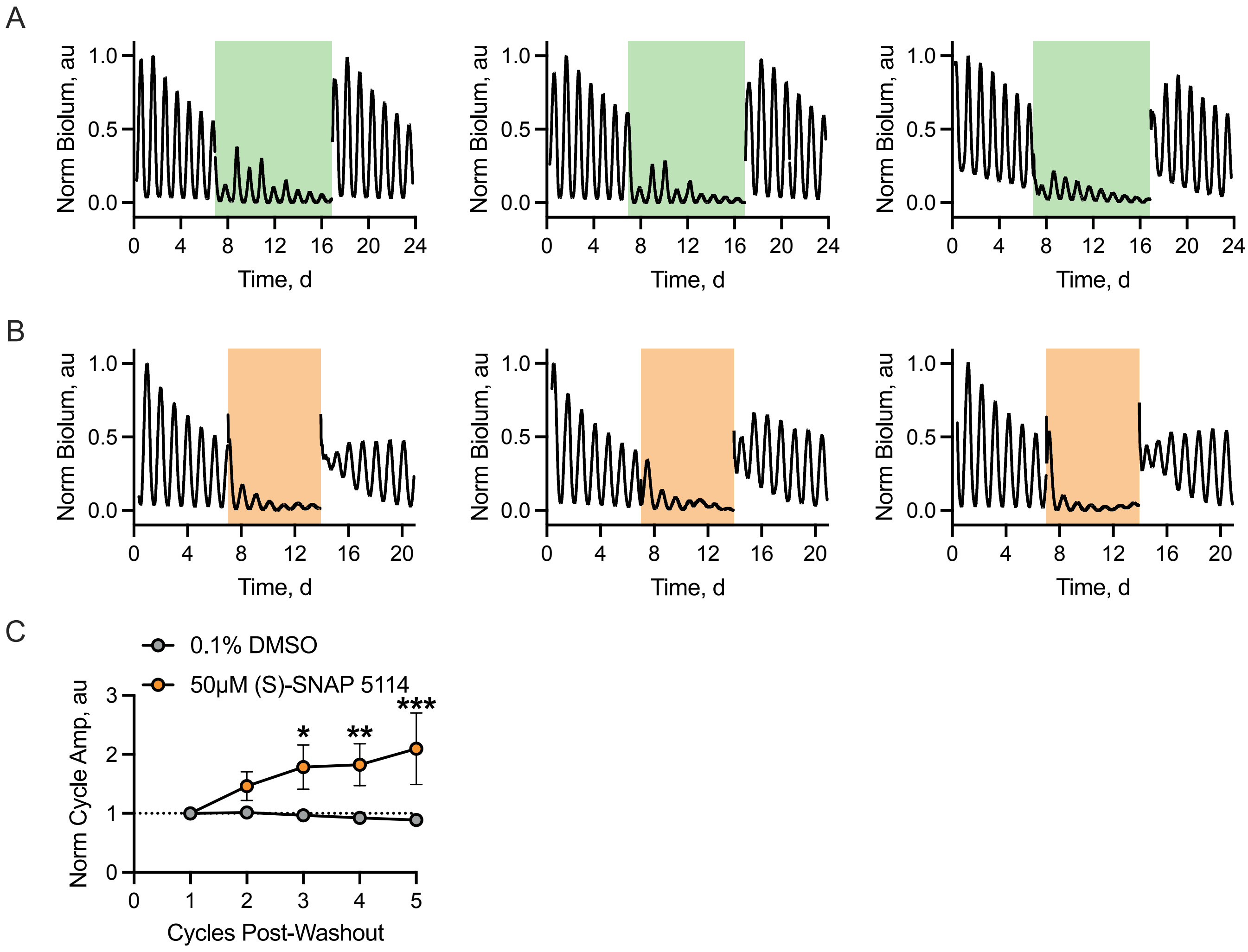

**Supplementary Figure 6: Observed and extrapolated peaks and troughs in iGABASnFR-rhythms under treatment with vehicle or (S)-SNAP 5114**

Example traces of raw fluorescence for slices treated with vehicle (upper, grey) or 50μM (S)-SNAP5114 (lower, orange). Predicted peak positions based on linear projections made from the baseline interval are shown as black circles, while predicted troughs are shown as maroon circles.

Supplementary Figure 6

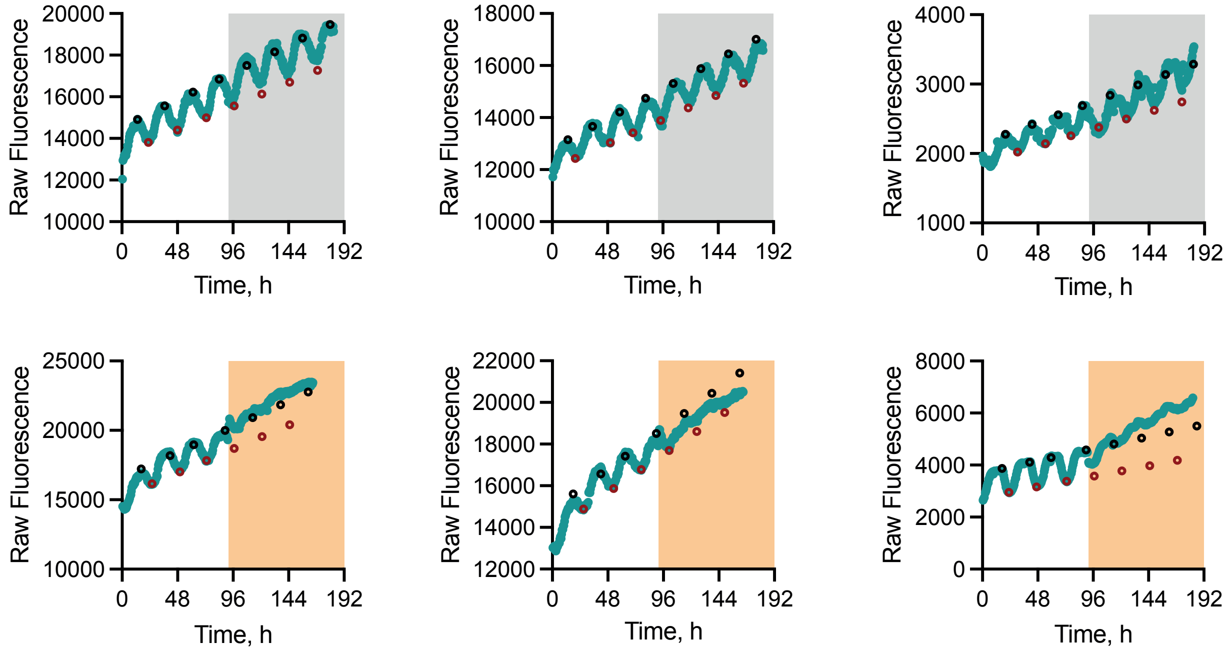

**Supplementary Figure 7: AlkK treatments that initiate rhythmicity in CRY1,2-null SCN are reversible**

A. Example traces of PER2::LUC bioluminescence from CRY1,2-null slices pre-treatment, during treatment and post-treatment with 10mM AlkK. Treatment interval is indicated by purple shading. B. Rhythmicity index calculated for baseline (BL), 10mM AlkK treatment (AlkK) and washout (WO) intervals. For B: N=11. Statistics: repeated-measures one-way ANOVA, $F(2,20)=15.5$, $p<0.0001$, Tukey's multiple comparisons test: ***$p<0.0006$ (comparison between BL and WO is non-significant, $p=0.88$).

Supplementary Figure 7

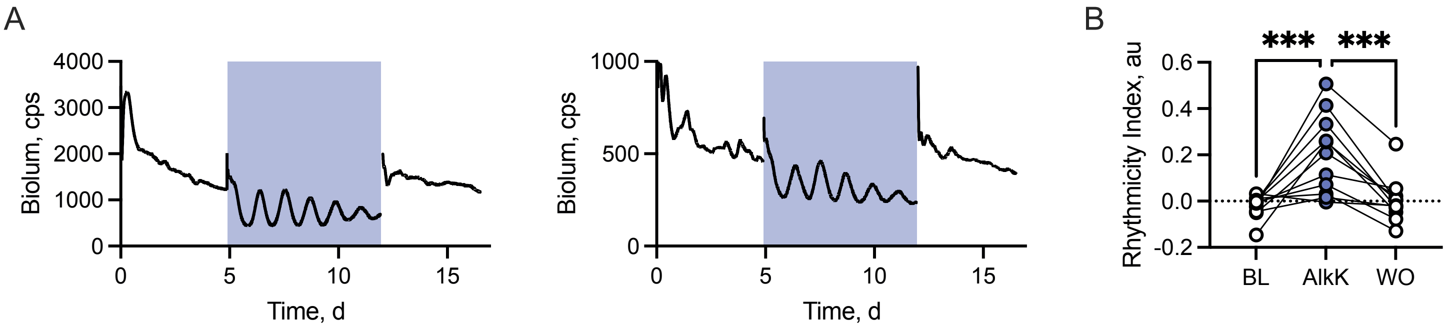

**Supplementary Methods**

**Seurat Script**

```
#Coarsely combine, cluster and identify by time and cell-type
(neurons, astrocytes, everything else)
#day-night SCN slice data from Morris et al, 2021 EMBO J to
query gene expression
#Version 1.1 written by Andrew Patton, 10th August 2022

#Load required packages

library(Seurat)
library(dplyr)
library(patchwork)
library(hdf5r)
library(ggplot2)

#Import raw data downloaded from accession database

mmSCN5.data <-
Read10X_h5('GSM5115763_mmSCN5_filtered_gene_bc_matrices_h5.h5'
)
mmSCN10.data <-
Read10X_h5('GSM5115764_mmSCN10_filtered_gene_bc_matrices_h5.h5
')
mmSCN6.data <-
Read10X_h5('GSM5115760_mmSCN6_filtered_gene_bc_matrices_h5.h5'
)
mmSCN7.data <-
Read10X_h5('GSM5115761_mmSCN7_filtered_gene_bc_matrices_h5.h5'
)
mmSCN11.data <-
Read10X_h5('GSM5115762_mmSCN11_filtered_gene_bc_matrices_h5.h5
')

#Create seurat objects from raw data

mmSCN5 <- CreateSeuratObject(counts = mmSCN5.data, project =
"mmSCN5", min.cells = 3, min.features = 200)
mmSCN10 <- CreateSeuratObject(counts = mmSCN10.data, project =
"mmSCN10", min.cells = 3, min.features = 200)
mmSCN6 <- CreateSeuratObject(counts = mmSCN6.data, project =
"mmSCN6", min.cells = 3, min.features = 200)
mmSCN7 <- CreateSeuratObject(counts = mmSCN7.data, project =
"mmSCN7", min.cells = 3, min.features = 200)
mmSCN11 <- CreateSeuratObject(counts = mmSCN11.data, project =
"mmSCN11", min.cells = 3, min.features = 200)

#Merge datasets irrespective of time of day for cell-type
clustering
```

```
All <- merge(mmSCN5, y = c(mmSCN10, mmSCN6, mmSCN7, mmSCN11),
add.cell.ids = c("SCN5", "SCN10", "SCN6", "SCN7", "SCN11"),
project = "All")
All
rm(mmSCN5.data, mmSCN10.data, mmSCN6.data, mmSCN7.data,
mmSCN11.data, mmSCN5, mmSCN6,mmSCN7, mmSCN10, mmSCN11)
gc()

#QC to remove mitochondrial genes

All[["percent.mt"]] <- PercentageFeatureSet(All, pattern =
"^mt-")
VlnPlot(All, features = c("nFeature_RNA", "nCount_RNA",
"percent.mt"), ncol = 3)
All <- subset(All, subset = nFeature_RNA > 100 & nFeature_RNA
< 8000 & percent.mt < 12.5)

#Normalise the data and run clustering algorithms

All <- NormalizeData(All, normalization.method =
"LogNormalize", scale.factor = 10000)
All <- FindVariableFeatures(All, selection.method = "vst",
nfeatures = 2000)
all.genes <- rownames(All)
All <- ScaleData(All, features = all.genes)
All <- RunPCA(All, features = VariableFeatures(object = All),
npcs=100)
All <- FindNeighbors(All, dims = 1:15)
All <- FindClusters(All, resolution = 0.05)
All <- RunUMAP(All, dims = 1:15)
DimPlot(All, reduction = "umap")

#Check markers and assign cluster identity coarsely to allow
targetted interrogation of neuronal or astrocytic cell groups

All.markers <- FindAllMarkers(All, only.pos = TRUE, min.pct =
0.25, logfc.threshold = 0.25)
VlnPlot(All, features = c("Tubb3", "Slc32a1", "Celf4","Gfap",
"Aldh1l1", "Aqp4", "Ndrg2", "S100b"))

new.cluster.ids <- c("Neurons",
"Neurons","Astrocytes","Other", "Other", "Astrocytes",
"Other", "Other", "Other")
names(new.cluster.ids) <- levels(All)
All <- RenameIdents(All, new.cluster.ids)
DimPlot(All, reduction = "umap", label = TRUE, pt.size = 0.5)
+ NoLegend()

saveRDS(All, file = "~/RProjects/Analysis/All.rds")

#Set up new metadata to allow backtracking to cell-type
clusters
```

```
All$CellType <- Idents(All)

#Switch active ident to the original idents to allow time of
day alignment

Idents(All) <- "orig.ident"
table(Idents(All))

#Subset the dataset to get daytime (CT7.5) or nighttime
(CT15.5) runs

All.Day <- subset(All, ident = c("mmSCN6","mmSCN7","mmSCN11"))
Idents(All.Day) <- "CellType"

All.Night <- subset(All, ident = c("mmSCN5","mmSCN10"))
Idents(All.Night) <- "CellType"

#Reassign cluster identities to take into account time cells
were harvested

new.cluster.ids.Day <- c("Neurons CT7.5", "Astrocytes CT7.5",
"Other CT7.5")
names(new.cluster.ids.Day) <- levels(All.Day)
All.Day <- RenameIdents(All.Day, new.cluster.ids.Day)

new.cluster.ids.Night <- c("Neurons CT15.5", "Astrocytes
CT15.5", "Other CT15.5")
names(new.cluster.ids.Night) <- levels(All.Night)
All.Night <- RenameIdents(All.Night, new.cluster.ids.Night)

#Merge time-aligned data sets back together and order the
cell-types

All.Time <- merge(All.Day, y= All.Night, merge.data = "TRUE",
project = "All.Time")
All.Time@active.ident <- factor(All.Time@active.ident,
levels=c("Neurons CT7.5", "Astrocytes CT7.5", "Other CT7.5",
"Neurons CT15.5", "Astrocytes CT15.5", "Other CT15.5"))

#Create dot plot to show expression of genes of interest
(GATs) alongside sanity check
#(Tubb3 [B-Tubulin, Neurons] and Aldh1l1 [Astrocytes])

DotPlot(All.Time, features = c("Slc6a1", "Slc6a13", "Slc6a11",
"Slc6a12", "Tubb3", "Aldh1l1")) + scale_color_viridis_c(option
= "magma")

saveRDS(All.Time, file = ~/RProjects/Analysis/AllTime.rds")
```

```
#Redo clustering analysis to restore order and allow further
future analyses - merge does not carry over previous
clustering results

All.Time.Umap <- All.Time
All.Time.Umap$CellTypeT <- Idents(All.Time.Umap)

All.Time.Umap <- NormalizeData(All.Time.Umap,
normalization.method = "LogNormalize", scale.factor = 10000)
All.Time.Umap <- FindVariableFeatures(All.Time.Umap,
selection.method = "vst", nfeatures = 2000)

all.time.umap.genes <- rownames(All.Time.Umap)
All.Time.Umap <- ScaleData(All.Time.Umap, features =
all.time.umap.genes)
All.Time.Umap <- RunPCA(All.Time.Umap, features =
VariableFeatures(object = All.Time.Umap), npcs=100)
All.Time.Umap <- FindNeighbors(All.Time.Umap, dims = 1:15)
All.Time.Umap <- FindClusters(All.Time.Umap, resolution =
0.05)
All.Time.Umap <- RunUMAP(All.Time.Umap, dims = 1:15)
DimPlot(All.Time.Umap, reduction = "umap")

#Reassign previous clusters

Idents(All.Time.Umap) <- "CellTypeT"
saveRDS(All.Time.Umap, file =
"~/RProjects/Analysis/AllTime.rds")

#Run differential expression

DvN.Astros <- FindMarkers(All.Time.Umap, ident.1 = "Astrocytes
CT7.5", ident.2 = "Astrocytes CT15.5", test.use = "wilcox")

#Recreate heatmap

DoHeatmap(All.Time.Umap, features = c("Slc6a1", "Slc6a13",
"Slc6a11", "Slc6a12", "Tubb3", "Aldh1l1")) +
scale_fill_continuous(type="viridis")
```
